# Understanding and predicting ligand efficacy in the mu-opioid receptor through quantitative dynamical analysis of complex structures

**DOI:** 10.1101/2024.01.20.576427

**Authors:** Gabriel Tiago Galdino, Olivier Mailhot, Rafael Najmanovich

**Author notes:** These authors contributed equally (Gabriel Tiago Galdino, Olivier Mailhot).

## Abstract

The *µ*-opioid receptor (MOR) is a G-protein coupled receptor involved in nociception and is the primary target of opioid drugs. Understanding the relationships between ligand structure, receptor dynamics, and efficacy in activating MOR is crucial for drug discovery and development. Here, we use coarse-grained normal mode analysis to predict ligand-induced changes in receptor dynamics with the Quantitative Dynamics Activity Relationships (QDAR) DynaSig-ML methodology, training a LASSO regression model on the entropic signatures (ES) computed from ligand-receptor complexes. We train and validate the methodology using a dataset of 179 MOR ligands with experimentally measured efficacies split into strickly chemically different cross-validation sets. By analyzing the coefficients of the ES LASSO model, we identified key residues involved in MOR activation, several of which have mutational data supporting their role in MOR activation. Additionally, we explored a contacts-only LASSO model based on ligand-protein interactions. While the model showed predictive power, it failed at predicting efficacy for ligands with low structural similarity to the training set, emphasizing the importance of receptor dynamics for predicting ligand-induced receptor activation. Moreover, the low computational cost of our approach, at 3 CPU seconds per ligand-receptor complex, opens the door to its application in large-scale virtual screening contexts. Our work contributes to a better understanding of dynamics-function relationships in the *µ*-opioid receptor and provides a framework for predicting ligand efficacy based on ligand-induced changes in receptor dynamics.

## 1 INTRODUCTION

Opioid receptors are a family of G-Protein Coupled Receptors (GPCRs) characterized by their pivotal role in mediating analgesia. The family is composed of the three main types *µ, κ*, and *δ* (MOR, KOR and DOR), *µ* being the primary target of morphine and most analgesic opioids [1]. Although a wide range of FDA-approved opioids are available, issues related to their common side effects, most notably addiction potential, have increased demand for either safer opioids or novel analgesics targeting alternative pathways [2]. This need also extends to the search of drugs with specific efficacy profiles such as antagonists, primarily used in the treatment of opioid intoxication and overdose, and partial agonists, ligands that partially activate the receptor (even at saturating concentrations), but can achieve the same therapeutic effects as agonists and are reported to have a higher therapeutic index (analgesia versus depression) [3].

Among the strategies used for opioid discovery, *in silico* studies have the advantage of searching large libraries of molecules at a reduced cost compared to completely experimental strategies [4]. In the case of opioids, virtual screening methodologies are a promising approach given the availability of high-resolution structures and the variety of known ligands [5]. Most of those methods, however, select molecules based on their predicted free energy of binding (docking score), and are thus completely agnostic to the potential efficacy of the molecule. Studies pursuing *in silico* functional selection only apply it to a limited number of drug candidates, which prevents the high-throughput computational search for ligands with specific functional outcomes (agonists, antagonists, or partial agonists) [6].

The activation of GPCRs primarily depends on the stabilization of one or more active forms of the receptor as a result of the binding of a ligand and/or an effector. Even though this model is still under discussion, it is irrefutable that receptor dynamics play a major role in activation and is crucial for the understanding of the function of GPCR ligands [7].

Some studies employing ligand similarity or contact fingerprints in combination with machine learning techniques to classify ligands as either agonists or antagonists showed the potential of employing machine learning to incorporate functional selection in virtual screening [8, 9, 10, 11, 12, 13]. However, those methodologies are based on static properties of ligands and/or ligand/receptor complexes and are consequently exclusively sensitive to interactions within the biding pocket. Thus, these models do not consider the allosteric effects of ligand binding, which are fundamental to signal transduction in GPCRs [7].

Previous attempts to use molecular dynamics (MD) simulations for ligand-GPCR complexes in different activation states showed that important mechanistic information can be obtained for both agonists and antagonists [14, 15]. Those techniques, though, are constrained by the substantial computational time needed to carry out simulations that provide enough observation time for the study of large domain movements [16]. This limitation hinders the ability to compare a wide range of ligands with varying measured efficacies, making it challenging to identify objective dynamics-based measures that can be used to develop generalizable ligand efficacy predictors.

Alternatively, normal mode analysis (NMA) is a fast analytical technique that can capture the underlying dynamics of the biomolecule under study, representing any configuration within the entire conformational space and solving motions at all timescales within a single solution. Depending on the degree of coarse-graining, NMA may require only a few CPU seconds [17] for modestly-sized proteins. The Elastic Network Contact Model (ENCoM) is a coarse-grained NMA model that is sensitive to the all-atom context of the input structure [18]. ENCoM has recently been extended to account for nucleic acids and small molecules in addition to proteins [19]. ENCoM has been employed to study microRNA maturation [20], the emergence of SARS-CoV-2 variants [21] as well as the determinants of thermophile protein stability [18], among others. The recently introduced DynaSig-ML Python package [22] facilitates the combination of ENCoM with simple machine learning models to capture quantitaty dynamics-activity relationships (QDAR) emerging from large datasets of experimental measures. In the present work, we use the DynaSig-ML methodology to study *µ*-opioid receptor activation as explained by ligand-induced changes in receptor dynamics. We also compare these dynamics-based predictions with corresponding models trained on molecular contacts alone, and find that the inclusion of dynamics greatly enhances the generalizability of the models to classes of ligands chemically distant from the ligands seen in training. Our results pave the way to the application of ultra-high-throughput prediction of ligand efficacy in future virtual screening campaigns against opioid receptors, with a low compute cost of 3 CPU-seconds per ligand-receptor complex. Moreover, the LASSO linear model trained on all available data identifies known features of opioid receptor activation, further validating our methodology.

## 2 RESULTS AND DISCUSSION

### 2.1 MOR ligands dataset overview

To investigate whether simple physicochemical properties of molecules can predict their activity, we calculated the molecular weight and cLogP values for all selected ligands (see subsection 3.1 and subsection 3.2). By comparing the distribution of both variables for Low Efficacy Ligands, ligands with *E*_*max*_ lower than 50% for [^35^S]GTP*γ*S binding assays [23] (LEL), and High Efficacy Ligands, ligands with *E*_*max*_ higher than 50% (HEL), we observe that HEL are slightly smaller and more hydrophilic than LEL (Supplementary Figure 1), with a mean MW of 414±75 for HEL and 435±70 for LEL, and a mean cLogP of 3.24±1.70 for HEL and 3.41±1.76 for LEL. The differences in MW and cLogP are not sufficient to differentiate between HEL and LEL and do not correlate with *E*_*max*_ (Pearson’s r *<* 0.1).

We clustered all ligands using complete linkage (threshold of 0.4 Tanimoto similarity coefficient) to create validation sets mimicking a virtual screening situation where a predictive model is applied to chemotypes outside what was seen in training. The complete linkage ensures that all ligands which are outside a given cluster have a similarity coefficient under 0.4 to any ligand inside the cluster, this threshold is less than a half from 0.85 (value typically applied for drug like molecules [24]). Figure 1 shows the resulting clustering and the class of each ligand. Two large clusters of over 60 molecules appear: one containing morphinan molecules, which we call morphine-like; the other containing molecules similar to fentanyl, called fentanyl-like. In addition to these two main clusters, five small clusters of 11 ligands or less are present. Within each cluster, further similarity between the ligands does not clearly separate HEL from LEL, with no apparent pattern emerging on the dendrogram. In terms of the *E*_*max*_ distributions, the morphine-like cluster has a flatter shape, whereas the fentanyl-like cluster has two distinct peaks at either ends of the spectrum. However, both are relatively well balanced in their proportions of HEL and LEL: the morphine-like cluster has 44.3% LEL while the fentanyl-like cluster has 55.2% LEL.

**Figure 1:**
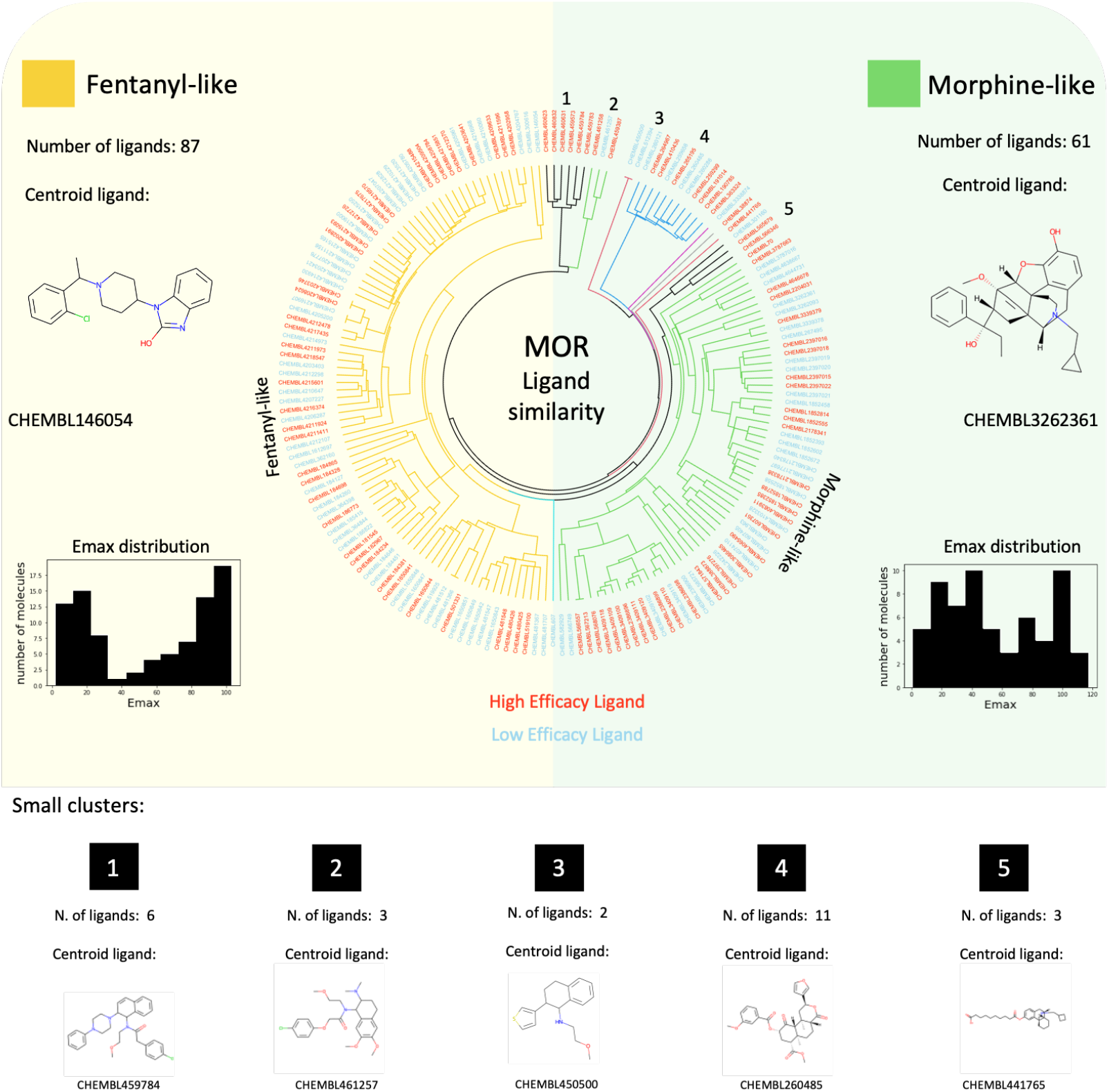
MOR ligands similarity clustering overview. The fan dendrogram **(center)** shows the clustering of all ligands based on their Tanimoto similarity coefficient derived using 1024-bit Morgan fingerprints with a radius of 2. Complete linkage clustering is performed with a similarity threshold of 0.4. The label colors represent the classification of each ligand (red: high efficacy ligands and cyan: low efficacy ligands). Two large clusters (fentanyl-like and morphine-like molecules) and 5 small clusters were identified. The total number of molecules and the centroid ligand are given for all clusters, as well as the *E*_*max*_ distribution for fentanyl-like **(right)** and morphine-like **(left)** ligands.

### 2.2 Docking score does not correlate with efficacy

We utilized the FlexAID docking software [25] to predict the complex structure for each ligand in complex with MOR in its active state. The FlexAID complementarity function (CF) scores molecular poses (arbitrary units of energy) such that a lower score represents a more favorable predicted molecular binding [25]. A total of 179 MOR ligands (95 HEL and 84 LEL) resulted in a successful docking, as defined by having at least 10 distinct docking poses with a negative CF score. Figure 2 presents the classification performance of the docking score alone for the 179 MOR ligands. In Figure 2B, the ROC curve and its AUC (0.41) are compared to the best performing random simulation (AUC of 0.63). The AUC value being under 0.5 is a strong indicator that the CF score alone fails in classifying ligands as HEL or LEL. Moreover, the distribution of random AUCs (Figure 2C) shows that the performance is indeed not statistically significant, with a p-value of 0.98. Inspecting Figure 2A, one could imagine that whilst the average CF for all poses is not a good classifier for Emax, a classifier trained instead on the best pose (pose with the lowest CF) could be useful. The performance for a predictor considering only the best CF value is presented in supplementary figure 5. The ROC-AUC in this case is very similar for the one considering the mean CF for the 10 poses (ROC-AUC: 0.40 and p-value:0.99). Considering the above results, we decided not to include the docking score as an additional predictor variable in subsequent regression models.

**Figure 2:**
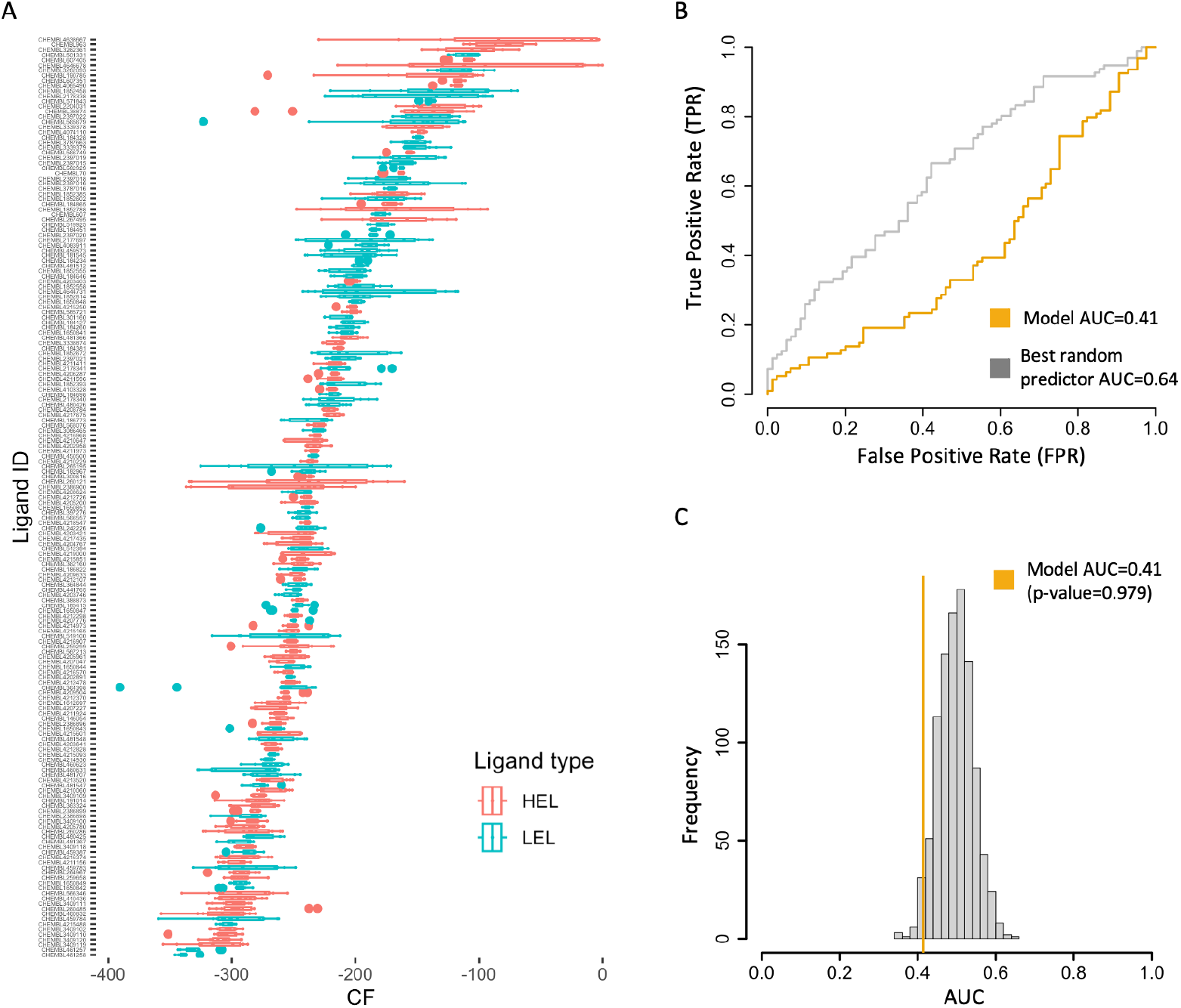
MOR docking simulation results. **(A)** Box-plot of FlexAID docking scores (CF) for the 10 poses of each ligand, ordered according to their mean CF value. The colors represent the classification of each ligand (red: High Efficacy Ligands and cyan: Low Efficacy Ligands). **(B)** ROC curve (AUC: 0.40) for CF as a classifier in comparison to the ROC curve of the best random predictor (AUC: 0.63). **(C)** Model AUC value in comparison to the distribution of the 1000 random predictors (p-value = 0.979)

It is important to highlight that the inability of the docking score to predict *E*_*max*_ is expected and doesn ‘t reflect inaccuracy in the scoring function. The absence of discriminatory power between LEL and HEL can be explained by the fact that ligand efficacy depends on relative ligand binding preference for specific receptor states (active, inactive, bound/unbound to the effector, and transition states between these) [26]. This relative preference of ligands for specific states over others is not reflected in the overall free energy of binding, which is the quantity estimated by the FlexAID scoring function.

### 2.3 Dynamical signatures-based LASSO regression model predicts ligand efficacy

We utilized DynaSig-ML [22] to calculate entropic signatures (ES) (a vector of entropic mean square fluctuations representing the flexibility of the complex) of all ligand-receptor complexes to test the predictive value of a dynamics-derived efficacy predictor (see subsection 3.4). The predictor was constructed using LASSO linear regression [27] and the perfomance was evaluated using two cross validation tests: leave-one-ligand-out (LOLO) and leave-one-cluster-out (LOCO) (see subsection 3.5). For both cross-validation tests, a set of 13 *β* scaling factor values from *e*^*−*3^ to *e*^3^ in log increments of 0.5 and a set of 25 regularization strengths (*α*) varying from 2^*−*3^ to 2^1^ in log_2_ increments of 0.25 were explored, for a total of 325 parameter combinations. The AUC under ROC for classification and Pearson linear correlation coefficient of all tested LASSO models for both validation tests are reported in the supplementary figure 2. For both LOLO and LOCO applied to the fentanyl-like cluster, the best performance is achieved when *β* = *e*^*−*0.5^. Additionally, LOCO applied to the morphine-like cluster achieves optimal performance when *β* = 1 (*e*^0^). Lower values of *α* (2^*−*3^ to 2^*−*2^) lead to slightly higher performance in LOLO, however LOCO benefits from higher regularization strengths, with optimum performance reached at *α* = 2^0.75^. This is consistent with LOCO cross-validation being a better test of the models ‘ generalizability: it is expected that higher regularization will drive coefficients of noisy variables down while keeping coefficients responsible for most of the signal.

Figure 3 shows the LOLO performance for *α* = 2^0.75^ and *β* = *e*^*−*0.5^, the predictor performs significantly better than random (AUC = 0.78, p-value: *<* 0.001 and person ‘s r= 0.48) and shows a very good enrichment of LEL with predicted *E*_*max*_ *<* 50% and HEL with predicted *E*_*max*_ *>* 50%. The performance of the model in the LOCO cross-validation test is shown in Figure 3. For both clusters, fentanyl-like and morphine-like, the model performs significantly better than random in classifying the ligands as high or low efficiency (ROC-AUC = 0.80 and 0.78, respectively, and p-values *<* 0.001). This performance illustrates that the model generalizes well to molecules structurally distant from the ones in the training set, namely with a Tanimoto similarity coefficient *<* 0.4. Thus, we hypothesize that such a model would hold its predictive value in a large-scale virtual screening scenario, where one is typically interested in finding diverse ligands topologically distant from known binders [28]. It is important to note that even though LOLO is an easier test then LOCO, the *α* value chosen was optimized for getting the best performance in LOCO. Also, LOLO has a performace similar to that in LOCO for the morphine like cluster, since in both cases the *E*_*max*_ distribuition of the molecules in the training set is equally distributed (Figure 1) making the classification problem harder, while in the fentanyl like cluster the molecules are either antagonists or full agonists.

**Figure 3:**
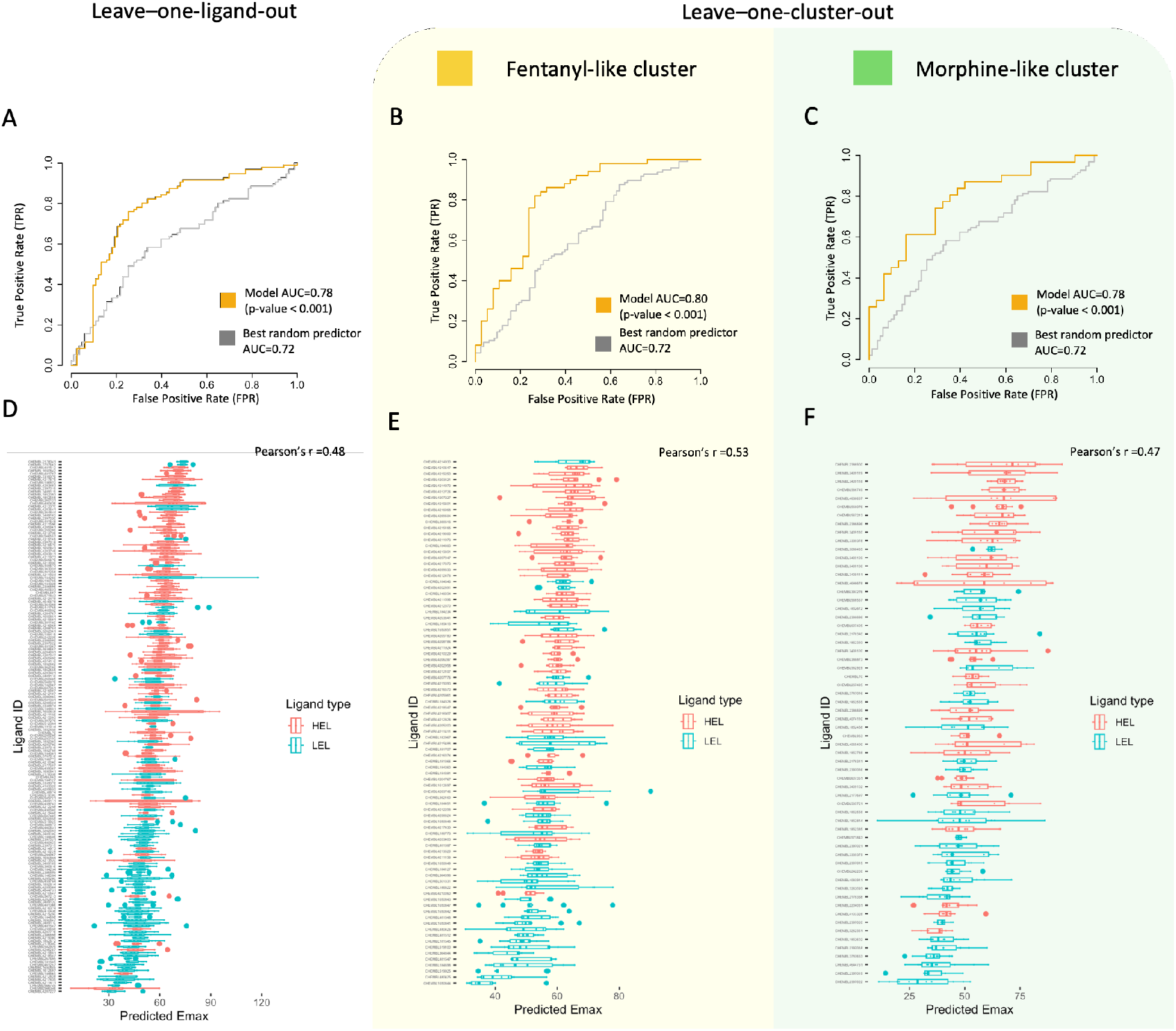
Validation of the Entropic Signature LASSO model. **(A-C)** ROC curves for leave-one-ligand-out (LOLO) and leave-one-cluster-out (LOCO) cross-validations, compared with the ROC curve from the best random predictor from 1000 replicates (equivalent to p=0.001). For LOCO (*α* = 2^0.75^ and *β* = *e*^*−*0.5^), the two >60 molecules clusters are in turn left out (fentanyl-like, *α* = 2^0.75^ and *β* = *e*^*−*0.5^, and morphine-like, *α* = 2^0.75^ and *β* = 1). **(D-F)** Boxplots of the predict *E*_*max*_ for all ligands ordered according their mean predicted *E*_*max*_ for LOLO and LOCO cross-validations.

LOCO cross-validation can be viewed as the ultimate test of the model’s generalizability to chemotypes beyond those seen in training. For both the morphine-like and fentanyl-like cluster, *α* = 2^0.75^ regularization strength led to the best performance. The best scaling factor for the morphine-like cluster was *β* = 1, while for the fentanyl-like *β* = *e*^*−*0.5^. We thus chose *α* = 2^0.75^ and *β* = *e*^*−*0.5^ as the parameters of a final LASSO model using entropic signatures as variables,this time fitting the model to all available data (179 MOR ligands, 10 docking poses each, see Methods). The resulting model would be a prime candidate for application to a prospective virtual screening campaign aiming at identifying novel MOR ligands with a given target *E*_*max*_ range. Owing to the model’s low computational cost (approximately 3 seconds CPU time per ligand-receptor complex), it could be applied to the full list of top docking hits (typically up to a million molecules [29]) for a modest computational cost. The predicted *E*_*max*_ values would then constitute an additional filter to apply before experimental testing, enriching ligands with the desired efficacy in the final set. Provided enough efficacy data are available for a wide diversity of known ligands, our approach could be applied to GPCRs other than MOR, and even other classes of targets where allostery plays a key role in the effect of ligand binding.

### 2.4 Structural and functional interpretation of LASSO coefficients

While DynaSig-ML can in principle utilize other machine learning classification methods, LASSO, as a linear regression technique, offers the unique advantage of interpretability of the results, allowing to isolate the contribution of unique residues and their mapping back on to the structure of the receptor. Figure 4A shows the coefficients for the final entropic signature LASSO model trained on all available data, using the parameters that led to the best generalizability on the LOCO cross-validations (*α* = 2^0.75^ and *β* = *e*^*−*0.5^). Negative coefficients mean that rigidification of that residue upon ligand binding leads to higher *E*_*max*_, while positive coefficients mean that softening of the position is correlated with higher *E*_*max*_. The *α* = 2^0.75^ regularization strength applied is sufficiently large to drive more than 85% of the LASSO coefficients to zero; remaining positions thus have the largest and most robust influence on receptor activation. A direct comparison between the coefficients ‘ magnitude is possible since the entropic signatures were standardized before training. The values of the LASSO coefficients are shown in Figure 4A and are are mapped onto the MOR structure in Figure 4B. This final entropic signature LASSO model is expected to be generalizable to novel opioid chemotypes and would be a prime candidate for application in a virtual screening campaign for the identification of ligands with specific *E*_*max*_ values. An interesting class of ligands to enrich would be partial agonists with predicted *E*_*max*_ around 60%, as these are expected to lead to analgesia with fewer side-effects and lower potential for addiction [3].

**Figure 4:**
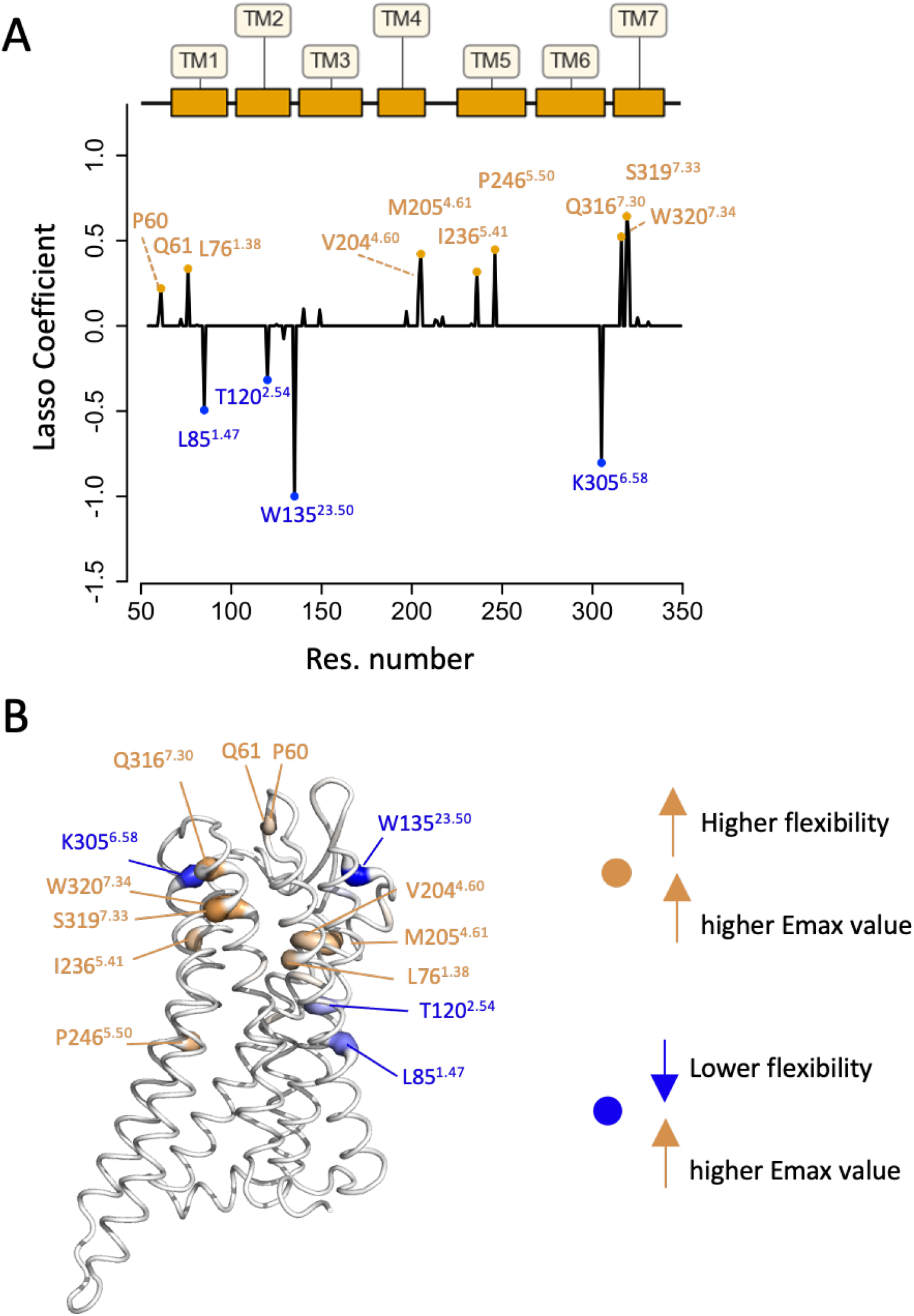
ES LASSO model trained with all available ligands. **(A)** LASSO coefficients at each receptor position along with a map of the seven transmembrane helices for reference. The 14 positions with highest absolute value coefficients are labeled. **(B)** LASSO coefficients mapped back to the MOR structure: the thickness of the cartoon represents the absolute value of the coefficient and the color represents the value with its sign (orange: positive, blue: negative and white/gray: neutral). Significant coefficients include:-0.33 at T120^2.54^ affecting tramadol response [30], -0.50 at L85^1.47^ with a variant (rs76773039) linked to morphine tolerance and dependence [30], 10.56 at W320^7.34^ influencing function and ligand bias [31], and -0.80 at K305^6.58^ important for selectivity in covalent binding of *β*-funaltrexamine (*β*-FNA) in the rat receptor [32].

The 14 positions with coefficients having an absolute value *>* 0.1 are labeled in Figure 4A-B. Positions *W*135^23.50^ and *K*305^6.58^ have coefficients of *−* 1 and*−*0.8, respectively, indicating that a rigidification of helix 6 and the extracellular loop 23 in the proximity of these positions leads to a higher *E*_*max*_. To the best of our knowledge, no variants in those positions have been reported to directly affect MOR activation. However, studies regarding the effect of *K*305^6.58^ on potency and the kinetics of certain ligands indicate that this residue plays an important role in activation [33, 32]. Two other positions, *L*85^1.47^ (coefficient: -0.5) and *T*120^2.54^ (coefficient:-0.3), have known variants listed in the ClinVar database [34]: L83I (code: rs76773039) and T120M (code:rs201310502). The mutation L83I is reported to affect morphine efficacy, increase receptor internalization, increase morphine tolerance, and is associated with higher morphine dependency [35], while T120M is reported to affect tramadol response [30].

Moreover, helices 5 and 7 seem to play an important role in activation by becoming more flexible around the binding pocket, as positions *Q*316^7.30^ (coefficient:0.5), *S*319^7.33^ (coefficient:0.7), *W*320^7.34^ (coefficient:0.5), *P*246^5.50^ (coefficient:0.4) and *I*236^5.41^ (coefficient:0.5) exhibit sizeable positive coefficients. Although mutagenesis data are not available for those specific residues in helix 7, Hothersall *et al*. [31] showed that the variant W320A in position *W*320^7.34^ has an effect on function and ligand bias. They also observed a clear uncoupling between mutation-driven changes in function and binding affinity for that position, a strong indication that changes in the dynamics in the region surrounding *W*320^7.34^ play an important role in activation. A list of all available variant data and all coefficients is provided in the supplementary material 1.

These findings illustrate that our model is effectively identifying relevant positions in terms of dynamics-function relationships of the *µ*-opioid receptor. It should be noted that our model is capturing changes in flexibility at the listed positions through propagation of ligand binding effects. Thus, it is not expected that positions identified as important would necessarily have functionally significant variants associated. However, the fact that several key positions in the model have reported functional impacts in the literature further validates our model and illuminates the crucial role of these residues in receptor activation.

### 2.5 Partial contact contribution model fails in predicting efficacy of novel ligands

The entropic signatures-based LASSO model significantly classifies MOR ligands as either high or low efficacy, even in the case of the LOCO cross-validations which present the model with ligands distant from ligands seen during training. However, since the underlying ENCoM model is based on surface area in contact [18], the predictive power could be coming mainly from a difference in atomic contacts within the binding pocket, as opposed to true dynamical effects. In order to disentangle the predictive power of dynamics and atomic contacts for efficacy prediction, we trained another set of LASSO models based on ligand-receptor atomic contacts. Importantly, the atom type definitions, surface in contact calculation and interaction matrix are identical for the contacts-based models and the entropic signatures-based models presented above (see subsection 3.3 and Supplementary Methods).

Figure 5A shows the performance of the contacts model for the LOLO cross-validation. The ROC-AUC of 0.73 is significantly better than random (p-value < 0.001), but worse than the performance of the entropic signatures (ES) model on the same test (ROC-AUC: 0.78, Figure 3A).

**Figure 5:**
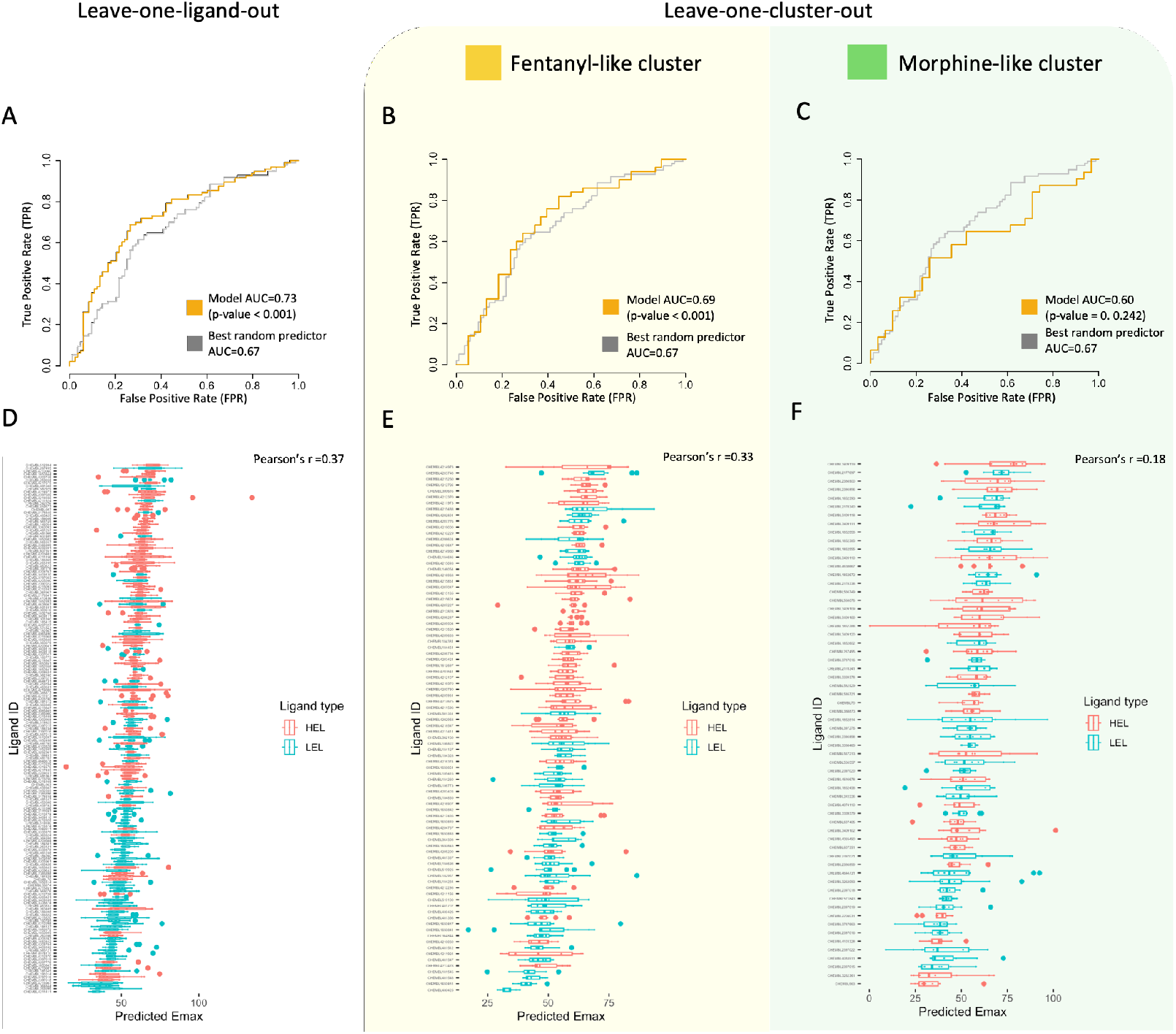
Validation of the atomic contacts LASSO model. **(A-C)** ROC curves for leave-one-ligand-out (LOLO) and leave-one-cluster-out (LOCO) cross-validations, compared with the ROC curve from the best random predictor from 1000 replicates (equivalent to p=0.001). For LOCO, the two >60 molecules clusters are in turn left out (fentanyl-like and morphine-like). **(D-F)** Boxplots of the predict *E*_*max*_ for all ligands ordered according their mean predicted *E*_*max*_ for LOLO and LOCO cross-validations.

On the two LOCO cross-validation tests (Figure 5B-C), the contacts model reaches 0.69 ROC-AUC (p-value < 0.001) for the fentanyl-like cluster of ligands and fails to reach statistical significance over a random classifier for the morphine-like cluster (ROC-AUC: 0.60, p-value = 0.242). Both values are substantially lower than what the ES model achieves, at 0.80 and 0.78 AUC-ROC, respectively (Figure 3B-C). We thus conclude that the contacts model fails to generalize when presented with novel chemotypes unseen during training. This failure is explained by the fact that ligands with different structures can achieve similar efficacies despite having different binding modes, as long as they affect the receptor ‘s allosteric networks in a similar fashion. Especially for GPCRs, the signal is propagated across the membrane, meaning that changes in dynamics are likely more robust for predicting efficacy than changes in binding site atomic contacts. This is indeed what our findings show: as stated before, the only additional information in the ES model is the context of receptor dynamics, as the changes in atomic contacts are modeled in identical ways within the ES and contacts model.

Despite the contacts model’s failure to generalize to novel chemotypes, it achieves statistically significant classification of the ligands in the LOLO cross-validation. In order to gain insight into what specific contacts hold predictive power for ligand efficacy, we trained a final contacts LASSO model using all available data, as we have done in Figure 4 (see Methods). To train as generalizable a contacts model as possible, we used *α* = 2^0.75^ regularization strength, as it led to the highest observed performance on the morphine-like LOCO cross-validation and a non-significant decrease in performance on the fentanyl-like cluster, as seen in Supplementary Figure 3.

Figure 6 shows the LASSO coefficients for the final contacts model. Positive coefficients mean that positive interaction energies between the residue in that position and the ligand (unfavorable contacts) lead to higher *E*_*max*_, while negative coefficients mean that favorable (negative) interactions with that position lead to higher *E*_*max*_. Figure 6A shows 12 coefficients with an absolute value higher than 0.1, and Figure 6B shows the coefficients mapped back to the structure. All 12 coefficients are from positions situated in the binding pocket and the majority of them are situated in helices 3 and 6. The residue *K*305^6.58^ is the only residue present in both the contacts and ES models, indicating its potential role as a key allosteric switch for the mu-opioid receptor. Additional residues identified here were previously identified as functionally relevant: Hothersall et al. [31] showed that mutations in *W*320^7.34^ lead to noticeable differences in function and affinity between agonists, highlighting its importance in receptor activation. *M*153^6.58^ is also predicted to be a ‘microswitch’ for some pathways in MOR activation [36].

**Figure 6:**
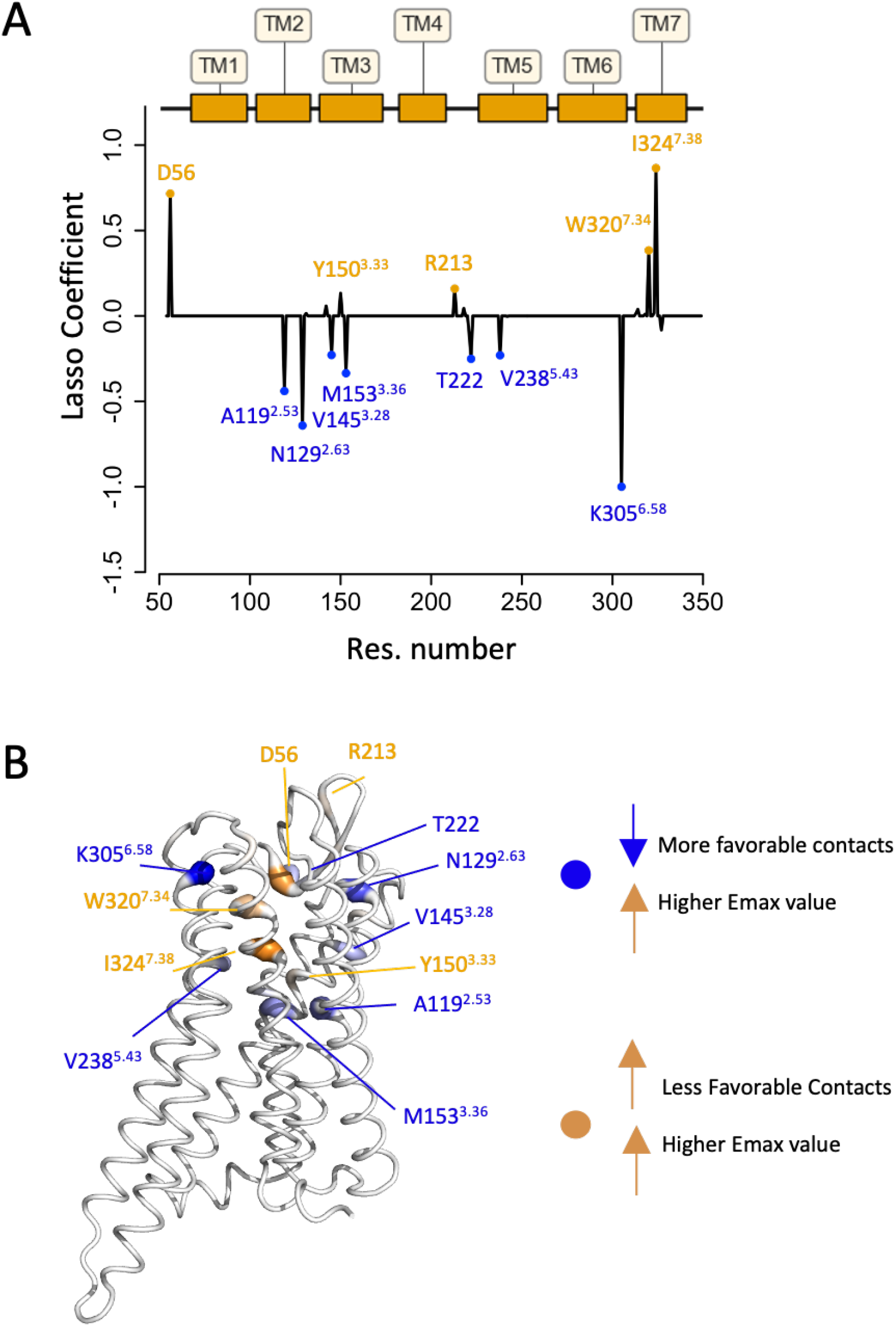
Contacts only LASSO model trained with all available ligands. **(A)** LASSO coefficients at each receptor position along with a map of the seven transmembrane helices for reference. The 12 positions with highest absolute value coefficients are labeled. **(B)** LASSO coefficients mapped back to the MOR structure: the thickness of the cartoon represents the absolute value of the coefficient and the color represents the value with its sign (orange: positive, blue: negative and white/gray: neutral). Notable coefficients include: 0.38 at W320^7.34^, associated with effects on function and ligand bias [31], -0.40 at M153^3.36^, known as a ‘microswitch’ for *β*-arrestin coupling for DAMGO and Fentanyl [36], and -1.00 at K305^6.58^, which is important for conferring selectivity for covalent binding of *β*-funaltrexamine (*β*-FNA) in the rat receptor [32].

Figure 7A shows the mean value of the partial CF contribution (variable in the contacts model) of each residue in the binding site of the receptor for every ligand in our dataset calculated with the Surfaces [37] software. The y-axis of the heat map is ordered based on the measured *E*_*max*_. The most striking pattern is that of residue *K*305^6.58^, showing unfavorable interactions with most of the LEL and either favorable or no interaction with HEL, as highlighted in Figure 7B. The docking poses of two typical ligands following the pattern are shown in Figure 7C. The *K*305^6.58^ residue is central in the binding pocket and has a contact in the poses of 140 out of the 179 ligands in our dataset. To evaluate if the mean partial CF for this residue can be used as an efficacy predictor, we constructed a ROC curve based on that single variable (Figure 7E). The AUC-ROC achieved by the *K*305^6.58^ mean partial CF is 0.73, statistically better than a random predictor at p < 0.001 (Figure 7F), but lower than the cross-validation performance of the ES LASSO model (AUC of 0.78-0.80).

**Figure 7:**
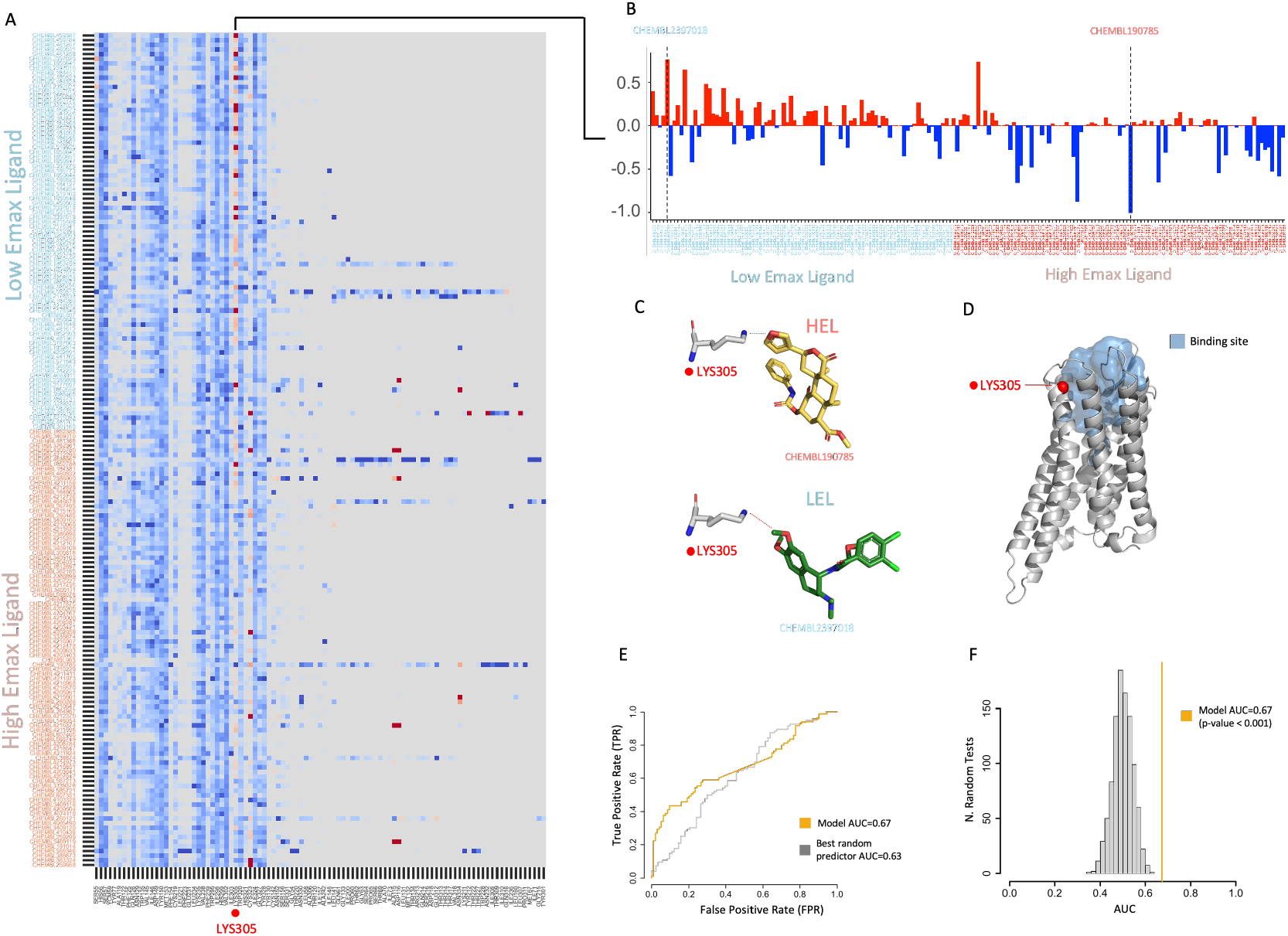
Contacts analysis of MOR Ligands. **(A)** Contact map showing the mean partial standardized complementarity function (CF) score for all residues in contact with at least one of the 10 poses of a ligand. **(B)** Focused view of contacts of all ligands with *K*305^6.58^. The height of each bar represents the color intensity in (A). **(C)** Two docking poses showing the interaction between *K*305^6.58^ and examples of high-efficacy and low-efficacy ligands. **(D)** MOR structure, the binding site area, and *K*305^6.58^. **(E)** ROC curve showing the performance of a classifier using the partial CF for *K*305^6.58^ (AUC=0.73). **(F)** AUC distribution for 1000 random predictors and p-value (<0.001) for the classifier using the partial CF of *K*305^6.58^.

An interesting comparison between the contacts and entropic signatures LASSO models is that of their coefficients mapped back to the MOR structure (Figure 4B and Figure 6B). The only common coefficient is that of *K*305^6.58^, highly negative for both models. In the case of the contacts model, this means that a strong interaction between the ligand and the lysine leads to higher *E*_*max*_; for the entropic signature model, the negative coefficient indicates that a rigidification of *K*305^6.58^ leads to higher *E*_*max*_. Both models thus learn an analogous role of lysine 305, since a stabilizing interaction leads to rigidification. An apparent difference between the ES and contacts models is that two coefficients for the ES model extend lower than the binding site towards the intracellular extremity of the receptor: the *P*246^5.50^ and *L*85^1.47^ residues. Such coefficients point again towards allostery being a large part of the ES model’s advantage over the contacts model. It should be mentioned that the absence of coefficients on more distal parts of the receptor for the ES model, such as the intracellular portions of TM6, does not imply these residues hold no predictive power. It is rather a feature of LASSO regression to drive most coefficients to zero when using high enough regularization strength. Moreover, if groups of variables are strongly correlated, LASSO will drop all but one from the group. Since the ES model uses the exact same atomic interaction energies between ligand and receptor as the contacts model, but reaches higher and more generalizable performance, it is safe to assume that it does capture allosteric features of ligand-induced MOR activation.

### 2.6 Availability

A Jupyter notebook is provided, allowing readers to train their own predictive ENCoM ES models with DynaSig-ML. The provided code will reproduce the ES LASSO model for LOCO and can be used as a template for users to create their own predictive models. One critical step in that process is the definition of atom types for every atom in every ligand. The atom types utilized are those used in FlexAID [25]. The code is available as a public depository in GitHub at https://github.com/NRGlab/MORDynaSig-ML.

### 2.7 Conclusions

We investigated the relationship between ligand-induced changes in *µ*-opioid receptor (MOR) dynamics and maximum efficacy (*E*_*max*_), as captured by the ENCoM coarse-grained normal mode analysis model using the DynaSig-ML Quantitative Dynamics Activity Relationship (QDAR) methodology. We docked a dataset of 179 diverse ligands with experimentally measured *E*_*max*_ and found that the docking score alone failed to accurately classify ligands as high or low efficacy. Ligands in our dataset showed some slight physicochemical differences between high efficacy ligands (HEL) and low efficacy ligands (LEL), but these differences did not correlate with efficacy.

To capture dynamics-function relationships, we trained entropic signatures (ES) LASSO models, using stringent leave-one-ligand-out (LOLO) and leave-one-cluster-out (LOCO) cross-validations. Despite the training and testing sets of LOCO cross-validations being chemically distant, the ES LASSO models capture significant signal, with an AUC-ROC of 0.78-0.80 and a Pearson ‘s r of 0.47-0.53. By analyzing the LASSO coefficients, we identified key residues involved in MOR activation. Positions such as *W*135^23.50^ and *K*305^6.58^ exhibited high negative coefficients, indicating that rigidification of these residues increased receptor activation. Mutational data supports the significance of these residues in MOR activation.

We also explored a contact-only LASSO model that considered the partial contribution of ligand-protein interactions to the docking score. While this model showed some predictive power, it importantly failed to predict efficacy for ligands with low structural similarity to the training set, emphasizing the limitations of models based on static properties in capturing receptor activation. Indeed, GPCRs transmit signal through allosteric networks which lead to sizeable conformational changes, such that it is unsurprising that a dynamics-based model outperforms a contacts-based one. Both models tested in our work use the exact same atomic energies for the ligand-receptor interaction, the difference being that these interactions energies are interpreted in the context of global receptor dynamics in the case of the entropic signature model. This shows how considering receptor dynamics for the prediction of GPCR ligand efficacy not only leads to higher accuracy, but better generalizability to novel chemotypes. This generalizability of the model, combined with its low computational cost of around 3 CPU seconds per ligand-receptor complex, make it very attractive for application to prospective virtual screening campaigns aimed at the discovery of novel ligands with efficacy within ranges of interest.

In conclusion, our study highlights the crucial role of ligand-induced changes in receptor dynamics in determining *µ*-opioid receptor ligands’ efficacies. The ES LASSO model, incorporating entropic dynamical signatures, outperformed the contacts-only model and provided valuable insights into key residues involved in receptor activation. This work contributes to a better understanding of the structure-function relationship of MOR and provides a framework for predicting ligand efficacy in prospective large-scale virtual screening campaigns.

## 3 methods

### 3.1 Data selection

We constructed a dataset ligands with reported efficacy for [^35^S]GTP*γ*S binding assay. This technique has been extensively used for the study of direct G-protein activation after ligand stimulation [23], such that hundreds of MOR ligands have [^35^S]GTP*γ*S measurements available in ChEMBL [38]. Despite the existence of more accurate techniques for the measurement of GPCR activation, [^35^S]GTP*γ*S is known for its sensitivity, while also being less susceptible to the effects of signal amplification when compared to methods that measure activity further down the signaling cascade [39].

All experimental data and SMILES codes for each ligand were obtained from the ChEMBL database [38]. We selected 288 MOR (ChEMBL ID CHEMBL233) ligands with measured *E*_*max*_ for [^35^S]GTP*γ*S binding assays [23] using DAMGO [40] as standard. We obtained initial 3D conformations for each ligand by using the Open Babel [41] API for Python, removing all small ions from the smiles code.

We separated the ligands in two classes: **High efficacy ligands (HEL)**: ligands with *E*_*max*_ higher than 50%, a class containing partial agonists with high *E*_*max*_ value and full agonists; **Low efficacy ligands (LEL)**: ligands with *E*_*max*_ lower than 50%, a class containing partial agonists with low *E*_*max*_ values and antagonists.

The molecular weight and the logP of each ligand were calculated using RDkit Python package [42].

### 3.2 Molecular docking

For the receptor structure, we chose the crystal structure of the active MOR bound to the agonist BU-72 and a G-protein-like nanobody, with a resolution of 2.1Å (PDB code: 5C1M) [43]. Despite this structure originating from a mouse receptor, we selected it based on three criteria:

1) sequence similarity: 98% to the human opioid receptor sequence (spanning residues 52-347);

2) high resolution: 0.8Å finer than the best-resolution structure available in the PDB for the active human MOR; and 3) completeness: no missing loops or residues.

We utilized Get-Cleft [44] to calculate the cavity of the reference ligand originally present in the crystal structure. Using the FlexAID docking software [25], we performed a self-docking of the reference ligand in order to establish the parameters for the docking of all ligands. From the self-docking we determined a minimum of 1000 generations, 1000 genes and a mutation rate of 0.05 to achieve an RSMD under 2.0 Å.

We then performed 50 docking simulations for each ligand using FlexAID. Each simulation was performed with 4000 generations, 4000 genes (16 times more energy evaluations than the minimum) and we kept the top 10 poses of each simulation (total of 500 poses per ligand, across 50 simulations). From the 288 ligands, 179 ligands were docked successfully, i.e. resulted in at least 10 unique poses with negative docking scores.

To generate robust validation sets and understand the chemical diversity of our dataset, we clustered the successful ligands according to their structure similarity. We computed the Morgan fingerprints with a radius of 2, on 1024 bits, and computed all pairwise Tanimoto similarity coefficients. We clustered the ligands using single-linkage clustering with a Tanimoto similarity cut-off of 0.4. This threshold was chosen so the molecules are divided in at least 2 big clusters. Also, the chosen cut-off is less than a half of the contested threshold of Tanimoto similarity of 0.85, normally adopted for drug-like molecules [24, **?**]. Using single-linkage ensures that all ligands outside a given cluster have aTanimoto similarity coefficient of at most 0.4 to any of the ligands within the cluster. The list of selected ligands is available in supplementary file 1.

### 3.3 Contacts analysis

We utilized the Surfaces [37] software to quantify the contribution of each contact between atoms of the ligand and residues in the binding pocket to each docking pose. We identified 98 residues with nonzero partial scores with at least one of the docking poses, across all ligands. We then constructed a vector for each ligand pose with the values of the residues with non-zero contact interaction and added to a data set that served as input to DynaSig-ML to train a LASSO model to predict *E*_max_ based on enthalpic contributions from residue contacts.

### 3.4 Computing dynamical signatures of ligand-receptor complexes with DynaSig-ML

We used a customized interaction matrix based on FlexAID [25] in the V4 term of the ENCoM potential (see Supplementary Methods) to ensure that the same pairwise atom-type pseudo-energetic interaction matrix used to dock the ligands and quantify contacts with Surfaces is used to define the spring constants used in the dynamical analyses. To prepare the ligands, we assigned FlexAID atom types for each ligand and we assigned the atom closest to the center of mass of the molecule as the position of the single node representing the ligand in the elastic network. For each of the 10 poses per ligand, we calculated entropic signatures (ES), as first introduced by Mailhot *et al*. [20]:

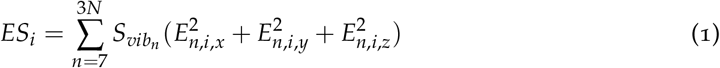

Where *ES*_*i*_ is the entropic mean square fluctuation for the *i*^*th*^ bead in the elastic network, N is the total number of beads, *E*_*n,i*_ is the movement of the bead in the *n*^*th*^ normal mode and *S*_*vibn*_ is the vibrational entropy associated to the *n*^*th*^ normal mode:

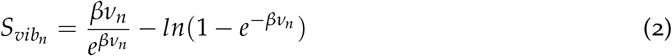

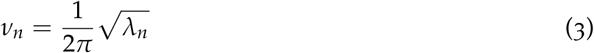

Where *v*_*n*_ is the vibrational frequency associated to eigenvalue of the *n*^*th*^ normal mode and *β* a thermodynamic scaling factor. We tested a set of 13 *β* varying between *e*^*−*3^ to *e*^3^ in log increments of 0.5. The ES were then standardized by position (flexibility factor for that position divided - mean flexibility factor for that position in all docking poses / standard deviation) before training the LASSO regression models.

### 3.5 LASSO linear regression

We performed LASSO linear regression [27] using either the ES or the contact vectors as the explanatory variables and the experimentally measured *E*_*max*_ as the outcome. We tested a set of 25 regularization strengths (*α*) varying from 2^*−*3^ to 2^1^ in log_2_ increments of 0.25.

Two validation strategies were employed: 1. **Leave-one-ligand-out (LOLO) cross-validation:** the 10 docking poses of one ligand constitute the testing set and all poses of the other ligands constitute the training set. The training and prediction procedure is repeated until all ligands have been tested, and the mean predicted values are pooled together and compared with measured *E*_*max*_ values for each combination of *β* and *α* values; 2. **Leave-one-cluster-out (LOCO) cross-validation:** all docking poses for all ligands within a large similarity cluster (cluster containing 60 or more ligands) are in the testing set and all other ligand poses are in the training set. This process is repeated for each large cluster and a list of predicted *E*_*max*_ for each combination of *β* and *α* values for all poses is registered for performance evaluation.

The performance evaluation was done based on two measures: Area Under the Curve (AUC) for the ROC curve constructed using the classification based on the measured *E*_*max*_ (HEL and LEL) and the predicted *E*_*max*_ of each molecule using sklearn [45]; and Pearson’s correlation r between the measured and predicted *E*_*max*_.

In order to calculate the significance (p-value) of each model performance, we constructed a distribution of 1000 replicates of N random *E*_*max*_ predictor models with the same mean and standard deviation as the distribution of measured *E*_*max*_ keeping the best AUC per replicate, where N below is the number of variables optimized that varies according to the model used, so that this simulations are equivalent to a Bonferroni correction [46]: ES model: 325 fitted linear regression models (13 *β* Χ 25 *α*) per replicate; Contacts model: 25 fitted linear regression models (25 *α*) per replicate; CF and single contact models: 1 fitted linear regression model per replicate.

### 3.6 Visualization of coefficients

To obtain a predictor trained with all ligands, we set aside 80% of the poses (8 poses per ligand) to be used as a training set and the remaining 20% served as a testing set. We trained a LASSO regression model on this dataset, with a single set of parameters chosen from the best performing parameters in the LOLO and LOCO cross-validation. The process was repeated 5 times, using every ligand pose in four out of the five LASSO models. For each position, the mean across the five trained LASSO models is taken as the coefficient of the final model. These LASSO coefficients for the contacts and ES models are plotted on the structure of MOR using PyMol [47]. The receptor is represented as a b-factor putty where the atoms are colored from blue (negative) to orange (positive) and the thickness of each position is related to the absolute value of that coefficient. For example, a residue with large negative coefficient indicates that a rigidification (in the case of the ES model) or a lower contact partial contribution (in the case of the contact model) in that position leads to higher *E*_*max*_.

## 4 SUPPLEMENTARY INFORMATION

File galdino2024 supplementary.pdf contains supplementary text, tables and figures. Supplementary Excel file galdino2024 supplementary data 1.xlsx contains the list of all ligands, the reference value of *E*_*max*_ obtained from CHEMBL, the cluster number and a list of all LASSO coefficients for all predictors and all variants found in for each position of MOR.

## REFERENCES

[1] William R Martin, W R Martin, C G Eades, J A Thompson, R E Huppler, and P E Gilbert. The effects of morphine- and nalorphine-like drugs in the nondependent and morphine-depenendent chronic spinal dog. Technical Report 3, 1976.

[2] Evan D. Kharasch, J. David Clark, and Jerome M. Adams. Opioids and Public Health: The Prescription Opioid Ecosystem and Need for Improved Management. Anesthesiology, 136(1):10–30, 1 2022.

[3] Joaquim Azevedo Neto, Anna Costanzini, Roberto De Giorgio, David G. Lambert, Chiara Ruzza, and Girolamo Calò. Biased versus partial agonism in the search for safer opioid analgesics. Molecules, 25, 9 2020.

[4] Giulio Poli, Marilisa Pia Dimmito, Adriano Mollica, Gokhan Zengin, Sandor Benyhe, Ferenc Zador, and Azzurra Stefanucci. Discovery of novel µ-opioid receptor inverse agonist from a combinatorial library of tetrapeptides through structure-based virtual screening. Molecules, 24(21), 10 2019.

[5] Albert J. Kooistra, Stefan Mordalski, Gáspár Pándy-Szekeres, Mauricio Esguerra, Alibek Mamyrbekov, Christian Munk, György M. Keseruű, and David E. Gloriam. GPCRdb in 2021: Integrating GPCR sequence, structure and function. Nucleic Acids Research, 49(D1):D335–D343, 1 2021.

[6] Hasan Pathan and John Williams. Basic opioid pharmacology: an update. British Journal of Pain, 6(1):11–16, 2 2012.

[7] H. Ongun Onaran and Tommaso Costa. Conceptual and experimental issues in biased agonism. Cellular Signalling, 82, 6 2021.

[8] Srilatha Sakamuru, Jinghua Zhao, Menghang Xia, Huixiao Hong, Anton Simeonov, Iosif Vaisman, and Ruili Huang. Predictive Models to Identify Small Molecule Activators and Inhibitors of Opioid Receptors, 6 2021.

[9] Elissa A. Fink, Jun Xu, Harald H übner, Joao M. Braz, Philipp Seemann, Charlotte Avet, Veronica Craik, Dorothee Weikert, Maximilian F. Schmidt, Chase M. Webb, Nataliya A. Tolmachova, Yurii S. Moroz, Xi Ping Huang, Chakrapani Kalyanaraman, Stefan Gahbauer, Geng Chen, Zheng Liu, Matthew P. Jacobson, John J. Irwin, Michel Bouvier, Yang Du, Brian K. Shoichet, Allan I. Basbaum, and Peter Gmeiner. Structure-based discovery of nonopioid analgesics acting through the α2A-adrenergic receptor. Science, 377(6614), 9 2022.

[10] Albert J. Kooistra, Rob Leurs, Iwan J.P. De Esch, and Chris De Graaf. Structure-based prediction of g-protein-coupled receptor ligand function: A β-adrenoceptor case study. Journal of Chemical Information and Modeling, 55(5):1045–1061, 5 2015.

[11] Jacob M. Remington, Kyle T. McKay, Noah B. Beckage, Jonathon B. Ferrell, Severin T. Schneebeli, and Jianing Li. GPCRLigNet: rapid screening for GPCR active ligands using machine learning. Journal of Computer-Aided Molecular Design, 37(3):147–156, 3 2023.

[12] Mireia Jim énez-Rosés, Bradley Angus Morgan, Maria Jimenez Sigstad, Thuy Duong Zoe Tran, Rohini Srivastava, Asuman Bunsuz, Leire Borrega-Román, Pattarin Hompluem, Sean A. Cullum, Clare R. Harwood, Eline J. Koers, David A. Sykes, Iain B. Styles, and Dmitry B. Veprintsev. Combined docking and machine learning identify key molecular determinants of ligand pharmacological activity on β2 adrenoceptor. Pharmacology Research and Perspectives, 10(5), 10 2022.

[13] Jooseong Oh, Hyi thaek Ceong, Dokyun Na, and Chungoo Park. A machine learning model for classifying G-protein-coupled receptors as agonists or antagonists. BMC Bioinformatics, 23, 8 2022.

[14] Marta Filizola, Kristen A Marino, and Yi Shang. Insights into the function of opioid receptors from molecular dynamics simulations of available crystal structures. British Journal of Pharmacology, 175:2834, 2018.

[15] Piotr F.J. Lipiński, Małgorzata Jarończyk, Jan Cz Dobrowolski, and Joanna Sadlej. Molecular dynamics of fentanyl bound to µ-opioid receptor. Journal of Molecular Modeling, 25(5), 5 2019.

[16] Brajesh Narayan, Ye Yuan, Arman Fathizadeh, Ron Elber, and Nicolae Viorel Buchete. Long-time methods for molecular dynamics simulations: Markov State Models and Milestoning. In Progress in Molecular Biology and Translational Science, volume 170, pages 215–237. Elsevier B.V., 1 2020.

[17] Jacob A. Bauer, Jelena Pavlovĺc, and Vladena Bauerov á-Hlinková. Normal mode analysis as a routine part of a structural investigation, 9 2019.

[18] Vincent Frappier and Rafael J. Najmanovich. A Coarse-Grained Elastic Network Atom Contact Model and Its Use in the Simulation of Protein Dynamics and the Prediction of the Effect of Mutations. PLoS Computational Biology, 10(4), 2014.

[19] Olivier Mailhot and Rafael Najmanovich. The NRGTEN Python package: an extensible toolkit for coarse-grained normal mode analysis of proteins, nucleic acids, small molecules and their complexes. Bioinformatics, 37(19):3369–3371, 10 2021.

[20] Olivier Mailhot, Vincent Frappier, François Major, and Rafael J. Najmanovich. Sequence-sensitive elastic network captures dynamical features necessary for miR-125a maturation. PLoS Computational Biology, 18(12), 12 2022.

[21] Natália Teruel, Olivier Mailhot, and Rafael J. Najmanovich. Modelling conformational state dynamics and its role on infection for SARS-CoV-2 Spike protein variants. PLoS Computational Biology, 17(8), 8 2021.

[22] Olivier Mailhot, François Major, and Rafael Najmanovich. The DynaSig-ML Python package: automated learning of biomolecular dynamics–function relationships. Bioinformatics, 39(4), 4 2023.

[23] C. Harrison and J. R. Traynor. The [35S]GTPγS binding assay: Approaches and applications in pharmacology. Life Sciences, 74(4):489–508, 12 2003.

[24] Yvonne C. Martin, James L. Kofron, and Linda M. Traphagen. Do structurally similar molecules have similar biological activity? Journal of Medicinal Chemistry, 45:4350–4358, 9 2002.

[25] Francis Gaudreault and Rafael J. Najmanovich. FlexAID: Revisiting Docking on Non-Native-Complex Structures. Journal of Chemical Information and Modeling, 55(7):1323–1336, 7 2015.

[26] Naomi R. Latorraca, A. J. Venkatakrishnan, and Ron O. Dror. GPCR dynamics: Structures in motion, 1 2017.

[27] Robert Tibshirani. Regression Shrinkage and Selection via the Lasso. Technical Report 1, 1996.

[28] Jiankun Lyu, Sheng Wang, Trent E Balius, Isha Singh, Anat Levit, Yurii S Moroz, Matthew J O‘Meara, Tao Che, Enkhjargal Algaa, Kateryna Tolmachova, et al. Ultra-large library docking for discovering new chemotypes. Nature, 566(7743):224–229, 2019.

[29] Brian J Bender, Stefan Gahbauer, Andreas Luttens, Jiankun Lyu, Chase M Webb, Reed M Stein, Elissa A Fink, Trent E Balius, Jens Carlsson, John J Irwin, et al. A practical guide to large-scale docking. Nature protocols, 16(10):4799–4832, 2021.

[30] S T Sherry, M.-H Ward, M Kholodov, J Baker, L Phan, E M Smigielski, and K Sirotkin. dbsnp: the ncbi database of genetic variation, 2001.

[31] J. Daniel Hothersall, Rubben Torella, Sian Humphreys, Monique Hooley, Alastair Brown, Gordon McMurray, and Sarah A. Nickolls. Residues W320 and Y328 within the binding site of the µ-opioid receptor influence opiate ligand bias. Neuropharmacology, 118:46–58, 5 2017.

[32] Kelly M. Dimattio, Chongguang Chen, Lei Shi, and Lee Yuan Liu-Chen. K3036.58 in the µ opioid (MOP) receptor is important in conferring selectivity for covalent binding of β-funaltrexamine (β-FNA). European Journal of Pharmacology, 748:93–100, 2 2015.

[33] Gregg Bonner, Fan Meng, and Huda Akil. Selectivity of m-opioid receptor determined by interfacial residues near third extracellular loop. Technical report, 2000.

[34] Melissa J. Landrum, Shanmuga Chitipiralla, Garth R. Brown, Chao Chen, Baoshan Gu, Jennifer Hart, Douglas Hoffman, Wonhee Jang, Kuljeet Kaur, Chunlei Liu, Vitaly Lyoshin, Zenith Maddipatla, Rama Maiti, Joseph Mitchell, Nuala O‘Leary, George R. Riley, Wenyao Shi, George Zhou, Valerie Schneider, Donna Maglott, J. Bradley Holmes, and Brandi L. Kattman. ClinVar: Improvements to accessing data. Nucleic Acids Research, 48(D1):D835–D844, 1 2020.

[35] Ajay Ravindranathan, Geoff Joslyn, Margaret Robertson, Marc A Schuckit, Jennifer L Whistler, and Raymond L White. Functional characterization of human variants of the mu-opioid receptor gene. Technical report.

[36] Parker W. de Waal, Jingjing Shi, Erli You, Xiaoxi Wang, Karsten Melcher, Yi Jiang, H. Eric Xu, and Bradley M. Dickson. Molecular mechanisms of fentanyl mediated Beta-arrestin biased signaling. PLoS Computational Biology, 16(4), 2020.

[37] Natália Teruel, Vinicius Magalhães Borges, and Rafael Najmanovich. Surfaces: a software to quantify and visualize interactions within and between proteins and ligands. Bioinformatics, 39, 10 2023.

[38] David Mendez, Anna Gaulton, A. Patrícia Bento, Jon Chambers, Marleen De Veij, Eloy F élix, María Paula Magariños, Juan F. Mosquera, Prudence Mutowo, Michał Nowotka, María Gordillo-Maranñón, Fiona Hunter, Laura Junco, Grace Mugumbate, Milagros Rodriguez-Lopez, Francis Atkinson, Nicolas Bosc, Chris J. Radoux, Aldo Segura-Cabrera, Anne Hersey, and Andrew R. Leach. ChEMBL: Towards direct deposition of bioassay data. Nucleic Acids Research, 47(D1):D930–D940, 1 2019.

[39] Ru Zhang and Xin Xie. Tools for GPCR drug discovery, 3 2012.

[40] Balraj K Handa, Anthony C Lane, John A H Lord, Barry A Morgan, Michael J Rance, Colin F C Smith, B K Handa, A C Lane, J A H Lord, B A Morgan, M J Rance, and C F C Smith. ANALOGUES OF ∼-LPHm.64 POSSESSING SELECTIVE AGONIST ACTIVITY AT/∼-OPIATE RECEPTORS. Technical report, 1981.

[41] Noel M. O‘Boyle, Michael Banck, Craig A. James, Chris Morley, Tim Vandermeersch, and Geoffrey R. Hutchison. Open Babel: An Open chemical toolbox. Journal of Cheminformatics, 3(10), 10 2011.

[42] Greg Landrum. Rdkit documentation. Release, 1:1–79, 2013.

[43] Weijiao Huang, Aashish Manglik, A. J. Venkatakrishnan, Toon Laeremans, Evan N. Feinberg, Adrian L. Sanborn, Hideaki E. Kato, Kathryn E. Livingston, Thor S. Thorsen, Ralf C. Kling, Sébastien Granier, Peter Gmeiner, Stephen M. Husbands, John R. Traynor, William I. Weis, Jan Steyaert, Ron O. Dror, and Brian K. Kobilka. Structural insights into µ-opioid receptor activation. Nature, 524(7565):315–321, 8 2015.

[44] Francis Gaudreault, Louis Philippe Morency, and Rafael J. Najmanovich. NRGsuite: A PyMOL plugin to perform docking simulations in real time using FlexAID. Bioinformatics, 31(23):3856–3858, 6 2015.

[45] Fabian Pedregosa Fabianpedregosa, Vincent Michel, Olivier Grisel Oliviergrisel, Mathieu Blondel, Peter Prettenhofer, Ron Weiss, Jake Vanderplas, David Cournapeau Fabian Pedregosa, G. ël Varoquaux, Alexandre Gramfort, Bertrand Thirion, Olivier Grisel, Vincent Dubourg, Alexandre Passos, Matthieu Brucher, Matthieu Perrot and Édouardand, and Édouard Duchesnay, and FR Édouard Duchesnay Edouardduchesnay. Scikit-learn: Machine Learning in Python Ga ël Varoquaux Bertrand Thirion Vincent Dubourg Alexandre Passos PEDREGOSA, VAROQUAUX, GRAMFORT ET AL. Matthieu Perrot. Technical report, 2011.

[46] Bonferroni Author and Francesco Brambilla. Statistica metodologica e calcolo delle probabilità (a proposito di recenti studi del prof. bonferroni), 1938.

[47] Warren L. DeLano. PyMOL User ‘s Guide. Technical report, DeLano Scientific LLC, 2004.

