## Supplementary material for "Understanding and predicting ligand efficacy in the mu-opioid receptor through quantitative dynamical analysis of complex structures": galdino2024_supplementary.pdf

[1]Department of Pharmacology and Physiology, Université de Montréal, Montreal, Canada
[2]Department of Pharmaceutical Chemistry, University of California, San Francisco, California, USA
[†]These authors contributed equally.
[*]To whom correspondence should be addressed.

January 20, 2024

Contact:

The original version of Elastic Network Contact Model (ENCoM) potential was first presented by *Frappier et al.* [1] It has a similar potential function to the one used in the Spring generalized Tensor Model (STeM) but includes a $\beta_{ij}$ factor that modulates the the non-bound interactions between two different atom types.

$$\begin{aligned} V_{\text{ENCoM}}(\vec{R}, \vec{R}_0) &= \sum_{\text{bonds}} V_1(r, r_0) + \sum_{\text{angles}} V_2(\theta, \theta_0) \\ &+ \sum_{\text{dihedrals}} V_3(\phi, \phi_0) + \sum_{i<j-3} V_4(r_{ij}, r_{ij_0}) \\ &= \sum_{\text{bonds}} \alpha_1 (r - r_0)^2 + \sum_{\text{angles}} \alpha_2 (\theta - \theta_0)^2 \\ &+ \sum_{\text{dihedrals}} \left[\alpha_3 (1 - \cos(\phi - \phi_0)) + \frac{\alpha_3}{2}(1 - \cos 3(\phi - \phi_0))\right] \\ &+ \sum_{i<j-3} (\beta_{ij} + \alpha_4) \left[5\left(\frac{r_{ij_0}}{r_{ij}}\right)^{12} - 6\left(\frac{r_{ij_0}}{r_{ij}}\right)^{10}\right] \end{aligned} \tag{1}$$

Those $\beta_{ij}$ factors are calculated from a set of different pairwise atom-type interactions ($\epsilon$) weighted by their surface in contact ($S$):

$$\beta_{ij} = \sum_{k}^{N_i} \sum_{l}^{N_j} \epsilon_{T(k)T(l)} S_{kl} \tag{2}$$

The default configuration of NRGten, heavy atoms are categorized into eight distinct types, following the classification system proposed by Sobolev et al. (1996). Within this framework, the parameter $\epsilon$ is assigned a binary value, which reflects whether the interaction is favorable or unfavorable:

$$\epsilon_{T(k)T(l)} = \begin{cases} \sigma_+ & \text{for favorable interactions} \\ \sigma_- & \text{for unfavorable interactions} \end{cases} \tag{3}$$

In this study, we opted to utilize the 40 atom-type definition delineated in FlexAID, assigning the parameter $\epsilon$ based on the interaction values determined by *Gaudreault et al.* [2]. Within the framework of FlexAID's Complementary Function (CF), interactions are considered more favorable as their values become more negative. Conversely, within the ENCoM potential context, this relationship is inverted, necessitating the multiplication of interaction values by -1. The specific values of each pairwise interaction utilized in our predictive model are detailed in Supplementary Table 1.

The ENCoM potential fundamentally represents a quadratic approximation of the potential energy, derived through a Taylor's series expansion. Consequently, repulsive values for the parameter $\beta_{ij}$ are not permitted. Therefore, in instances where the cumulative non-bonded interactions between any two nodes within the elastic network model yield a negative sum, the value of $\beta_{ij}$ is set to zero.

$$\beta_{ij} = \begin{cases} \sum_{k}^{N_i} \sum_{l}^{N_j} \epsilon_{T(k)T(l)} S_{kl}, & \text{if } \sum_{k}^{N_i} \sum_{l}^{N_j} \epsilon_{T(k)T(l)} S_{kl} > 0 \\ 0, & \text{otherwise} \end{cases} \tag{4}$$

### Table 1. 40 atom-type interaction matrix adapted from FlexAID

| 0 | 1 | 2 | 3 | 4 | 5 | 6 | 7 | 8 | 9 | 10 | 11 | 12 | 13 | 14 | 15 | 16 | 17 | 18 | 19 | 20 |
|---|---|---|---|---|---|---|---|---|---|---|---|---|---|---|---|---|---|---|---|---|
| 1 | 0.00 | 119.40 | 162.90 | 179.10 | 0.00 | 0.00 | 0.00 | 0.00 | 0.00 | -157.10 | -60.54 | 78.63 | 198.30 | 180.80 | -93.26 | 0.00 | 0.00 | 0.00 | 0.00 | 0.00 |
| 2 | 119.40 | 198.90 | 69.11 | 149.40 | -194.10 | -165.40 | 88.78 | 0.00 | -36.97 | -15.81 | 21.59 | 61.99 | 10.06 | -61.29 | -14.76 | 0.00 | 115.00 | 123.10 | -81.35 | 0.00 |
| 3 | 162.90 | 69.11 | 37.75 | 53.32 | 191.90 | 71.64 | 68.71 | 0.00 | -87.28 | 63.97 | -26.90 | 0.95 | 50.23 | 44.36 | 30.77 | 0.00 | 0.00 | 48.67 | 121.80 | 0.00 |
| 4 | 179.10 | 149.40 | 53.32 | 58.92 | 183.90 | 0.00 | 124.20 | 0.00 | -36.65 | 18.30 | -23.98 | 16.94 | 5.61 | 21.65 | -25.60 | 0.00 | 135.60 | 51.64 | -100.30 | 0.00 |
| 5 | 0.00 | -194.10 | 191.90 | 183.90 | 0.00 | 0.00 | 161.80 | 0.00 | -174.80 | 118.30 | -180.90 | 83.10 | -195.90 | -67.99 | 54.19 | 0.00 | 0.00 | 167.90 | 0.00 | 0.00 |
| 6 | 0.00 | -165.40 | 71.64 | 0.00 | 0.00 | 0.00 | 0.00 | 0.00 | 0.00 | 0.00 | 168.90 | 0.00 | 0.00 | 71.74 | 0.00 | 0.00 | 0.00 | 0.00 | 0.00 | 0.00 |
| 7 | 0.00 | 88.78 | 68.71 | 124.20 | 161.80 | 0.00 | 0.00 | 0.00 | -84.57 | 55.80 | 195.70 | 122.30 | 87.38 | 144.80 | 197.60 | 0.00 | 0.00 | -28.13 | 0.00 | 0.00 |
| 8 | 0.00 | 0.00 | 0.00 | 0.00 | 0.00 | 0.00 | 0.00 | 0.00 | 0.00 | 0.00 | 0.00 | 0.00 | 0.00 | 0.00 | 0.00 | 0.00 | 0.00 | 0.00 | 0.00 | 0.00 |
| 9 | 0.00 | -36.97 | -87.28 | -36.65 | -174.80 | 0.00 | -84.57 | 0.00 | -81.66 | 16.35 | -160.20 | -160.40 | 103.90 | 101.80 | 159.00 | 0.00 | 0.00 | 66.40 | 0.00 | 0.00 |
| 10 | -157.10 | -15.81 | 63.97 | 18.30 | 118.30 | 0.00 | 55.80 | 0.00 | 16.35 | -156.70 | 157.30 | 26.35 | 86.64 | 189.60 | 125.30 | 0.00 | 0.00 | 190.10 | 169.00 | 0.00 |
| 11 | -60.54 | 21.59 | -26.90 | -23.98 | -180.90 | 168.90 | 195.70 | 0.00 | -160.20 | 157.30 | -72.28 | -86.55 | 141.10 | 70.05 | 140.10 | 0.00 | 0.00 | -46.22 | -73.66 | 0.00 |
| 12 | 78.63 | 61.99 | 0.95 | 16.94 | 83.10 | 0.00 | 122.30 | 0.00 | -160.40 | 26.35 | -86.55 | -133.00 | 81.10 | 49.12 | 127.70 | 0.00 | 0.00 | 50.59 | -159.90 | 0.00 |
| 13 | 198.30 | 10.06 | 50.23 | 5.61 | -195.90 | 0.00 | 87.38 | 0.00 | 103.90 | 86.64 | 141.10 | 81.10 | -56.58 | 33.00 | -52.00 | 0.00 | 0.00 | -27.66 | -170.40 | 0.00 |
| 14 | 180.80 | -61.29 | 44.36 | 21.65 | -67.99 | 71.74 | 144.80 | 0.00 | 101.80 | 189.60 | 70.05 | 49.12 | 33.00 | 77.28 | 95.08 | 0.00 | 0.00 | 79.64 | -170.80 | 0.00 |
| 15 | -93.26 | -14.76 | 30.77 | -25.60 | 54.19 | 0.00 | 197.60 | 0.00 | 159.00 | 125.30 | 140.10 | 127.70 | -52.00 | 95.08 | -45.17 | 0.00 | 0.00 | -32.66 | 77.56 | 0.00 |
| 16 | 0.00 | 0.00 | 0.00 | 0.00 | 0.00 | 0.00 | 0.00 | 0.00 | 0.00 | 0.00 | 0.00 | 0.00 | 0.00 | 0.00 | 0.00 | 0.00 | 0.00 | 0.00 | 0.00 | 0.00 |
| 17 | 0.00 | 115.00 | 0.00 | 135.60 | 0.00 | 0.00 | 0.00 | 0.00 | 0.00 | 0.00 | 0.00 | 0.00 | 0.00 | 0.00 | 0.00 | 0.00 | 0.00 | 0.00 | 0.00 | 0.00 |
| 18 | 0.00 | 123.10 | 48.67 | 51.64 | 167.90 | 0.00 | -28.13 | 0.00 | 66.40 | 190.10 | -46.22 | 50.59 | -27.66 | 79.64 | -32.66 | 0.00 | 0.00 | 145.50 | -103.90 | 0.00 |
| 19 | 0.00 | -81.35 | 121.80 | -100.30 | 0.00 | 0.00 | 0.00 | 0.00 | 0.00 | 169.00 | -73.66 | -159.90 | -170.40 | -170.80 | 77.56 | 0.00 | 0.00 | -103.90 | 0.00 | 0.00 |
| 20 | 0.00 | 0.00 | 0.00 | 0.00 | 0.00 | 0.00 | 0.00 | 0.00 | 0.00 | 0.00 | 0.00 | 0.00 | 0.00 | 0.00 | 0.00 | 0.00 | 0.00 | 0.00 | 0.00 | 0.00 |
| 21 | 0.00 | 0.00 | 0.00 | 0.00 | 0.00 | 0.00 | 0.00 | 0.00 | 0.00 | 0.00 | 0.00 | 0.00 | 0.00 | 0.00 | 0.00 | 0.00 | 0.00 | 0.00 | 0.00 | 0.00 |
| 22 | 0.00 | -76.92 | 8.39 | 58.99 | -167.70 | 0.00 | 0.00 | 0.00 | -143.00 | 148.80 | 117.90 | -44.25 | -61.46 | 163.40 | -101.40 | 0.00 | 0.00 | 41.28 | 0.00 | 0.00 |
| 23 | 0.00 | 186.10 | 74.04 | 57.06 | 28.17 | 0.00 | 0.00 | 0.00 | 0.00 | -7.70 | 46.59 | 28.99 | -51.70 | 149.50 | -187.50 | 0.00 | 0.00 | -5.70 | 0.00 | 0.00 |
| 24 | 0.00 | 193.70 | 93.41 | 25.05 | 0.00 | 0.00 | 0.00 | 0.00 | -45.57 | -88.72 | -36.68 | 0.00 | -19.20 | 0.00 | 69.11 | 0.00 | 0.00 | 167.50 | 0.00 | 0.00 |
| 25 | 0.00 | 0.00 | 0.00 | 12.72 | 190.10 | 0.00 | 0.00 | 0.00 | -50.17 | -137.30 | 0.00 | 1.93 | 0.00 | 0.00 | 0.00 | 0.00 | 0.00 | 119.70 | 0.00 | 0.00 |
| 26 | 0.00 | -13.04 | 157.60 | 0.00 | 0.00 | 0.00 | 0.00 | 0.00 | 0.00 | 0.00 | 0.00 | 0.00 | -9.61 | 0.00 | 0.00 | 0.00 | 0.00 | 0.00 | 0.00 | 0.00 |
| 27 | 0.00 | 0.00 | 0.00 | 0.00 | 0.00 | 0.00 | 0.00 | 0.00 | 0.00 | 0.00 | 0.00 | 0.00 | 0.00 | 0.00 | 0.00 | 0.00 | 0.00 | 0.00 | 0.00 | 0.00 |
| 28 | 0.00 | -152.50 | -131.20 | 0.00 | 0.00 | 0.00 | 0.00 | 0.00 | 0.00 | 0.00 | 61.63 | -92.97 | 61.55 | -67.59 | 194.70 | 0.00 | 0.00 | 0.00 | 0.00 | 0.00 |
| 29 | 0.00 | 0.00 | 0.00 | 0.00 | 0.00 | 0.00 | 0.00 | 0.00 | 0.00 | 0.00 | 0.00 | 0.00 | 0.00 | -71.70 | 0.00 | 0.00 | 0.00 | 0.00 | 0.00 | 0.00 |
| 30 | 0.00 | 0.00 | 180.80 | 0.00 | 0.00 | 0.00 | 0.00 | 0.00 | 0.00 | 0.00 | 0.00 | 0.00 | 0.00 | 0.00 | 0.00 | 0.00 | 0.00 | 0.00 | 0.00 | 0.00 |
| 31 | 0.00 | -119.80 | 0.00 | 0.00 | 0.00 | 0.00 | 0.00 | 0.00 | 0.00 | 0.00 | 0.00 | -37.17 | 0.00 | 0.00 | 0.00 | 0.00 | 0.00 | 0.00 | 0.00 | 0.00 |
| 32 | 0.00 | 0.00 | 0.00 | 0.00 | 0.00 | 0.00 | 0.00 | 0.00 | 0.00 | 0.00 | 0.00 | 0.00 | 0.00 | 0.00 | 0.00 | 0.00 | 0.00 | 0.00 | 0.00 | 0.00 |
| 33 | 0.00 | -121.20 | 0.00 | 0.00 | 0.00 | 0.00 | 0.00 | 0.00 | 0.00 | 0.00 | 0.00 | 0.00 | 0.00 | 0.00 | 0.00 | 0.00 | 0.00 | 0.00 | 0.00 | 0.00 |
| 34 | 0.00 | 0.00 | 0.00 | 0.00 | 0.00 | 0.00 | 0.00 | 0.00 | 0.00 | 0.00 | 0.00 | 0.00 | 0.00 | 0.00 | 0.00 | 0.00 | 0.00 | 0.00 | 0.00 | 0.00 |
| 35 | 0.00 | 40.54 | -95.36 | -160.80 | 0.00 | 0.00 | 0.00 | 0.00 | -158.90 | 0.00 | 185.80 | 0.00 | 182.10 | 186.70 | 173.40 | 0.00 | 0.00 | 153.70 | 134.40 | 0.00 |
| 36 | 0.00 | 0.00 | 0.00 | 0.00 | 0.00 | 0.00 | 0.00 | 0.00 | 0.00 | 0.00 | 0.00 | 0.00 | 0.00 | 197.80 | 68.34 | 0.00 | 0.00 | 0.00 | 0.00 | 0.00 |
| 37 | 0.00 | 0.00 | 0.00 | 0.00 | 0.00 | 0.00 | 0.00 | 0.00 | 0.00 | 0.00 | 0.00 | 0.00 | 0.00 | 0.00 | 0.00 | 0.00 | 0.00 | 0.00 | 0.00 | 0.00 |
| 38 | 0.00 | 0.00 | 0.00 | 0.00 | 0.00 | 0.00 | 0.00 | 0.00 | 0.00 | 0.00 | 0.00 | 0.00 | 0.00 | 0.00 | 0.00 | 0.00 | 0.00 | 0.00 | 0.00 | 0.00 |
| 39 | 0.00 | 0.00 | 0.00 | 0.00 | 0.00 | 0.00 | 0.00 | 0.00 | 0.00 | 0.00 | 0.00 | 0.00 | 0.00 | 0.00 | 0.00 | 0.00 | 0.00 | 0.00 | 0.00 | 0.00 |
| 40 | -159.10 | -61.75 | -72.50 | -62.17 | -120.30 | -167.20 | -136.40 | 0.00 | -147.10 | -198.30 | -190.50 | -162.80 | -33.99 | -43.24 | -29.56 | 0.00 | 0.00 | -94.93 | 176.20 | 0.00 |

| 0 | 21 | 22 | 23 | 24 | 25 | 26 | 27 | 28 | 29 | 30 | 31 | 32 | 33 | 34 | 35 | 36 | 37 | 38 | 39 | 40 |
|---|---|---|---|---|---|---|---|---|---|---|---|---|---|---|---|---|---|---|---|---|
| 1 | 0.00 | 0.00 | 0.00 | 0.00 | 0.00 | 0.00 | 0.00 | 0.00 | 0.00 | 0.00 | 0.00 | 0.00 | 0.00 | 0.00 | 0.00 | 0.00 | 0.00 | 0.00 | 0.00 | -159.10 |
| 2 | 0.00 | -76.92 | 186.10 | 193.70 | 0.00 | -13.04 | 0.00 | -152.50 | 0.00 | 0.00 | -119.80 | 0.00 | -121.20 | 0.00 | 40.54 | 0.00 | 0.00 | 0.00 | 0.00 | -61.75 |
| 3 | 0.00 | 8.39 | 74.04 | 93.41 | 0.00 | 157.60 | 0.00 | -131.20 | 0.00 | 180.80 | 0.00 | 0.00 | 0.00 | 0.00 | -95.36 | 0.00 | 0.00 | 0.00 | 0.00 | -72.50 |
| 4 | 0.00 | 58.99 | 57.06 | 25.05 | 12.72 | 0.00 | 0.00 | 0.00 | 0.00 | 0.00 | 0.00 | 0.00 | 0.00 | 0.00 | -160.80 | 0.00 | 0.00 | 0.00 | 0.00 | -62.17 |
| 5 | 0.00 | -167.70 | 28.17 | 0.00 | 190.10 | 0.00 | 0.00 | 0.00 | 0.00 | 0.00 | 0.00 | 0.00 | 0.00 | 0.00 | 0.00 | 0.00 | 0.00 | 0.00 | 0.00 | -120.30 |
| 6 | 0.00 | 0.00 | 0.00 | 0.00 | 0.00 | 0.00 | 0.00 | 0.00 | 0.00 | 0.00 | 0.00 | 0.00 | 0.00 | 0.00 | 0.00 | 0.00 | 0.00 | 0.00 | 0.00 | -167.20 |
| 7 | 0.00 | 0.00 | 0.00 | 0.00 | 0.00 | 0.00 | 0.00 | 0.00 | 0.00 | 0.00 | 0.00 | 0.00 | 0.00 | 0.00 | 0.00 | 0.00 | 0.00 | 0.00 | 0.00 | -136.40 |
| 8 | 0.00 | 0.00 | 0.00 | 0.00 | 0.00 | 0.00 | 0.00 | 0.00 | 0.00 | 0.00 | 0.00 | 0.00 | 0.00 | 0.00 | 0.00 | 0.00 | 0.00 | 0.00 | 0.00 | 0.00 |
| 9 | 0.00 | -143.00 | 0.00 | -45.57 | -50.17 | 0.00 | 0.00 | 0.00 | 0.00 | 0.00 | 0.00 | 0.00 | 0.00 | 0.00 | -158.90 | 0.00 | 0.00 | 0.00 | 0.00 | -147.10 |
| 10 | 0.00 | 148.80 | -7.70 | -88.72 | -137.30 | 0.00 | 0.00 | 0.00 | 0.00 | 0.00 | 0.00 | 0.00 | 0.00 | 0.00 | 0.00 | 0.00 | 0.00 | 0.00 | 0.00 | -198.30 |
| 11 | 0.00 | 117.90 | 46.59 | -36.68 | 0.00 | 0.00 | 0.00 | 61.63 | 0.00 | 0.00 | 0.00 | 0.00 | 0.00 | 0.00 | 185.80 | 0.00 | 0.00 | 0.00 | 0.00 | -190.50 |
| 12 | 0.00 | -44.25 | 28.99 | 0.00 | 1.93 | 0.00 | 0.00 | -92.97 | 0.00 | 0.00 | -37.17 | 0.00 | 0.00 | 0.00 | 0.00 | 0.00 | 0.00 | 0.00 | 0.00 | -162.80 |
| 13 | 0.00 | -61.46 | -51.70 | -19.20 | 0.00 | -9.61 | 0.00 | 61.55 | 0.00 | 0.00 | 0.00 | 0.00 | 0.00 | 0.00 | 182.10 | 0.00 | 0.00 | 0.00 | 0.00 | -33.99 |
| 14 | 0.00 | 163.40 | 149.50 | 0.00 | 0.00 | 0.00 | 0.00 | -67.59 | -71.70 | 0.00 | 0.00 | 0.00 | 0.00 | 0.00 | 186.70 | 197.80 | 0.00 | 0.00 | 0.00 | -43.24 |
| 15 | 0.00 | -101.40 | -187.50 | 69.11 | 0.00 | 0.00 | 0.00 | 194.70 | 0.00 | 0.00 | 0.00 | 0.00 | 0.00 | 0.00 | 173.40 | 68.34 | 0.00 | 0.00 | 0.00 | -29.56 |
| 16 | 0.00 | 0.00 | 0.00 | 0.00 | 0.00 | 0.00 | 0.00 | 0.00 | 0.00 | 0.00 | 0.00 | 0.00 | 0.00 | 0.00 | 0.00 | 0.00 | 0.00 | 0.00 | 0.00 | 0.00 |
| 17 | 0.00 | 0.00 | 0.00 | 0.00 | 0.00 | 0.00 | 0.00 | 0.00 | 0.00 | 0.00 | 0.00 | 0.00 | 0.00 | 0.00 | 0.00 | 0.00 | 0.00 | 0.00 | 0.00 | 0.00 |
| 18 | 0.00 | 41.28 | -5.70 | 167.50 | 119.70 | 0.00 | 0.00 | 0.00 | 0.00 | 0.00 | 0.00 | 0.00 | 0.00 | 0.00 | 153.70 | 0.00 | 0.00 | 0.00 | 0.00 | -94.93 |
| 19 | 0.00 | 0.00 | 0.00 | 0.00 | 0.00 | 0.00 | 0.00 | 0.00 | 0.00 | 0.00 | 0.00 | 0.00 | 0.00 | 0.00 | 134.40 | 0.00 | 0.00 | 0.00 | 0.00 | 176.20 |
| 20 | 0.00 | 0.00 | 0.00 | 0.00 | 0.00 | 0.00 | 0.00 | 0.00 | 0.00 | 0.00 | 0.00 | 0.00 | 0.00 | 0.00 | 0.00 | 0.00 | 0.00 | 0.00 | 0.00 | 0.00 |
| 21 | 0.00 | 0.00 | 0.00 | 0.00 | 0.00 | 0.00 | 0.00 | 0.00 | 0.00 | 0.00 | 0.00 | 0.00 | 0.00 | 0.00 | 0.00 | 0.00 | 0.00 | 0.00 | 0.00 | 0.00 |
| 22 | 0.00 | 0.00 | 0.00 | 0.00 | 0.00 | 0.00 | 0.00 | 163.00 | 0.00 | 0.00 | 0.00 | 0.00 | 0.00 | 0.00 | -89.37 | 0.00 | 0.00 | 0.00 | 0.00 | -134.70 |
| 23 | 0.00 | 0.00 | 0.00 | 0.00 | 0.00 | 0.00 | 0.00 | 0.00 | 0.00 | 0.00 | 0.00 | 0.00 | 0.00 | 0.00 | 118.70 | 0.00 | 0.00 | 0.00 | 0.00 | -3.12 |
| 24 | 0.00 | 0.00 | 0.00 | 0.00 | 0.00 | 0.00 | 0.00 | 0.00 | 0.00 | 0.00 | 0.00 | 0.00 | 0.00 | 0.00 | 0.00 | 0.00 | 0.00 | 0.00 | 0.00 | -198.80 |
| 25 | 0.00 | 0.00 | 0.00 | 0.00 | 0.00 | 0.00 | 0.00 | 0.00 | 0.00 | 0.00 | 0.00 | 0.00 | 0.00 | 0.00 | 0.00 | 0.00 | 0.00 | 0.00 | 0.00 | -138.40 |
| 26 | 0.00 | 0.00 | 0.00 | 0.00 | 0.00 | 0.00 | 0.00 | 0.00 | 0.00 | 0.00 | 0.00 | 0.00 | 0.00 | 0.00 | 0.00 | 0.00 | 0.00 | 0.00 | 0.00 | 0.00 |
| 27 | 0.00 | 0.00 | 0.00 | 0.00 | 0.00 | 0.00 | 0.00 | 0.00 | 0.00 | 0.00 | 0.00 | 0.00 | 0.00 | 0.00 | 0.00 | 0.00 | 0.00 | 0.00 | 0.00 | 0.00 |
| 28 | 0.00 | 163.00 | 0.00 | 0.00 | 0.00 | 0.00 | 0.00 | 0.00 | 0.00 | 0.00 | 0.00 | 0.00 | 0.00 | 0.00 | 0.00 | 0.00 | 0.00 | 0.00 | 0.00 | 0.00 |
| 29 | 0.00 | 0.00 | 0.00 | 0.00 | 0.00 | 0.00 | 0.00 | 0.00 | 0.00 | 0.00 | 0.00 | 0.00 | 0.00 | 0.00 | 0.00 | 0.00 | 0.00 | 0.00 | 0.00 | 0.00 |
| 30 | 0.00 | 0.00 | 0.00 | 0.00 | 0.00 | 0.00 | 0.00 | 0.00 | 0.00 | 0.00 | 0.00 | 0.00 | 0.00 | 0.00 | 0.00 | 0.00 | 0.00 | 0.00 | 0.00 | 0.00 |
| 31 | 0.00 | 0.00 | 0.00 | 0.00 | 0.00 | 0.00 | 0.00 | 0.00 | 0.00 | 0.00 | 0.00 | 0.00 | 0.00 | 0.00 | 0.00 | 0.00 | 0.00 | 0.00 | 0.00 | 0.00 |
| 32 | 0.00 | 0.00 | 0.00 | 0.00 | 0.00 | 0.00 | 0.00 | 0.00 | 0.00 | 0.00 | 0.00 | 0.00 | 0.00 | 0.00 | 0.00 | 0.00 | 0.00 | 0.00 | 0.00 | 0.00 |
| 33 | 0.00 | 0.00 | 0.00 | 0.00 | 0.00 | 0.00 | 0.00 | 0.00 | 0.00 | 0.00 | 0.00 | 0.00 | 0.00 | 0.00 | 0.00 | 0.00 | 0.00 | 0.00 | 0.00 | 0.00 |
| 34 | 0.00 | 0.00 | 0.00 | 0.00 | 0.00 | 0.00 | 0.00 | 0.00 | 0.00 | 0.00 | 0.00 | 0.00 | 0.00 | 0.00 | 0.00 | 0.00 | 0.00 | 0.00 | 0.00 | 0.00 |
| 35 | 0.00 | -89.37 | 118.70 | 0.00 | 0.00 | 0.00 | 0.00 | 0.00 | 0.00 | 0.00 | 0.00 | 0.00 | 0.00 | 0.00 | 0.00 | 0.00 | 0.00 | 0.00 | 0.00 | 0.00 |
| 36 | 0.00 | 0.00 | 0.00 | 0.00 | 0.00 | 0.00 | 0.00 | 0.00 | 0.00 | 0.00 | 0.00 | 0.00 | 0.00 | 0.00 | 0.00 | 0.00 | 0.00 | 0.00 | 0.00 | 0.00 |
| 37 | 0.00 | 0.00 | 0.00 | 0.00 | 0.00 | 0.00 | 0.00 | 0.00 | 0.00 | 0.00 | 0.00 | 0.00 | 0.00 | 0.00 | 0.00 | 0.00 | 0.00 | 0.00 | 0.00 | 0.00 |
| 38 | 0.00 | 0.00 | 0.00 | 0.00 | 0.00 | 0.00 | 0.00 | 0.00 | 0.00 | 0.00 | 0.00 | 0.00 | 0.00 | 0.00 | 0.00 | 0.00 | 0.00 | 0.00 | 0.00 | 0.00 |
| 39 | 0.00 | 0.00 | 0.00 | 0.00 | 0.00 | 0.00 | 0.00 | 0.00 | 0.00 | 0.00 | 0.00 | 0.00 | 0.00 | 0.00 | 0.00 | 0.00 | 0.00 | 0.00 | 0.00 | 0.00 |
| 40 | 0.00 | -134.70 | -3.12 | -198.80 | -138.40 | 0.00 | 0.00 | 0.00 | 0.00 | 0.00 | 0.00 | 0.00 | 0.00 | 0.00 | 0.00 | 0.00 | 0.00 | 0.00 | 0.00 | 0.00 |

Values of $\epsilon$ defined accord to the ones proposed by *Gaudreault et al.* [2] The values were inverted in order to make favorable interactions positive (red) and unfavorable interactions negative (blue).

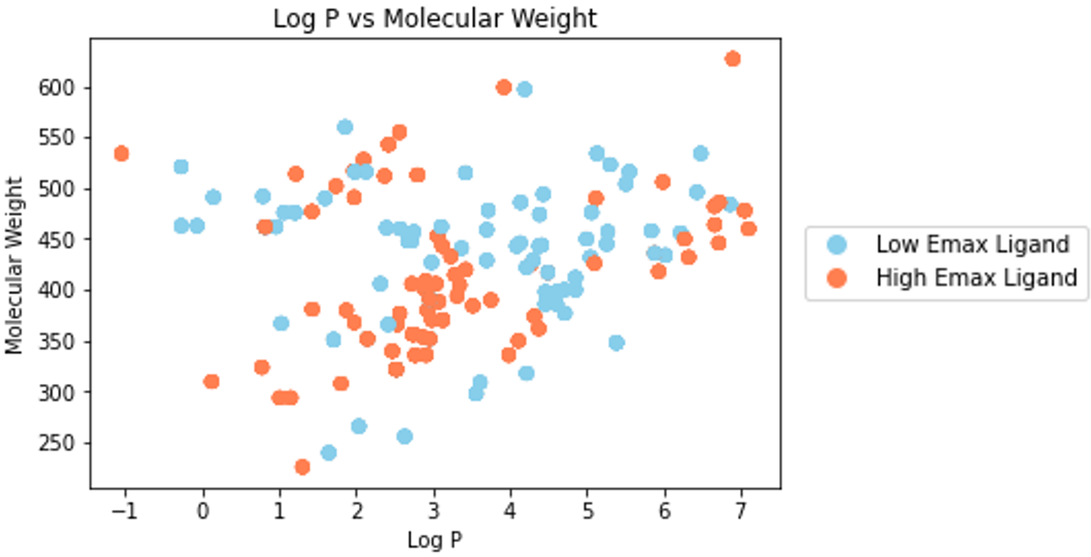


**Figure 1: Molecular Weight and log P values for all MOR ligands classified according their activity.**

### 3 ES MODEL PERFORMANCE OVERVIEW

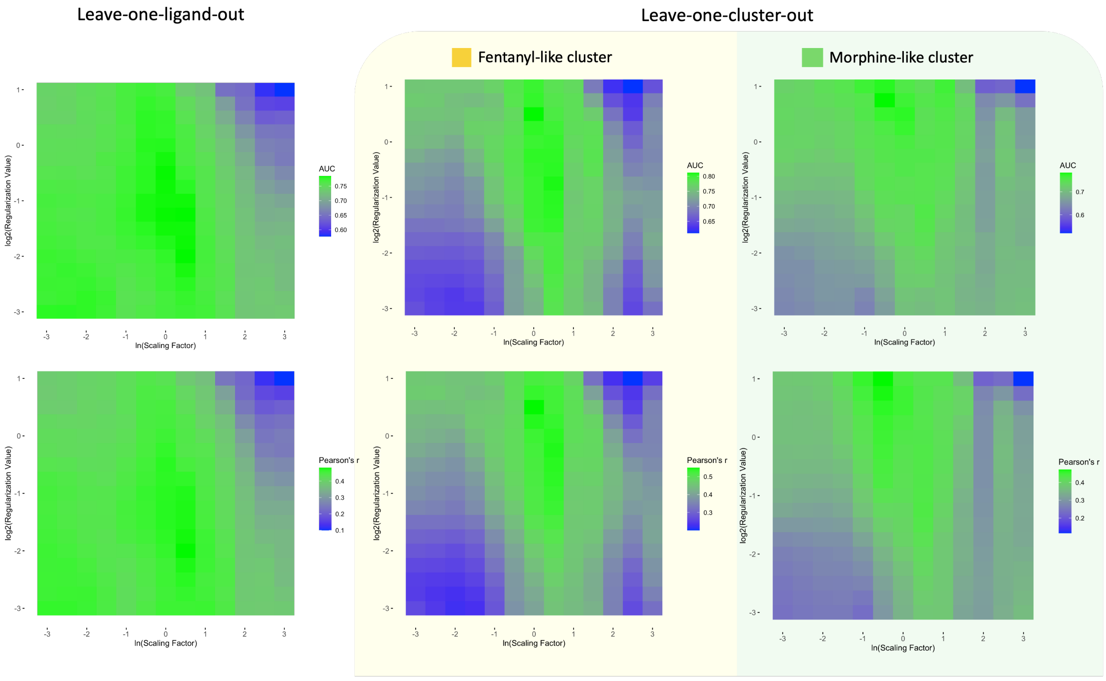


**Figure 2: Leave-one-ligand-out and leave-one-cluster-out performance (AUC and Pearson's R) over all combinations of Scaling Factors ($\beta$) and LASSO Regularization Values ($\alpha$).**

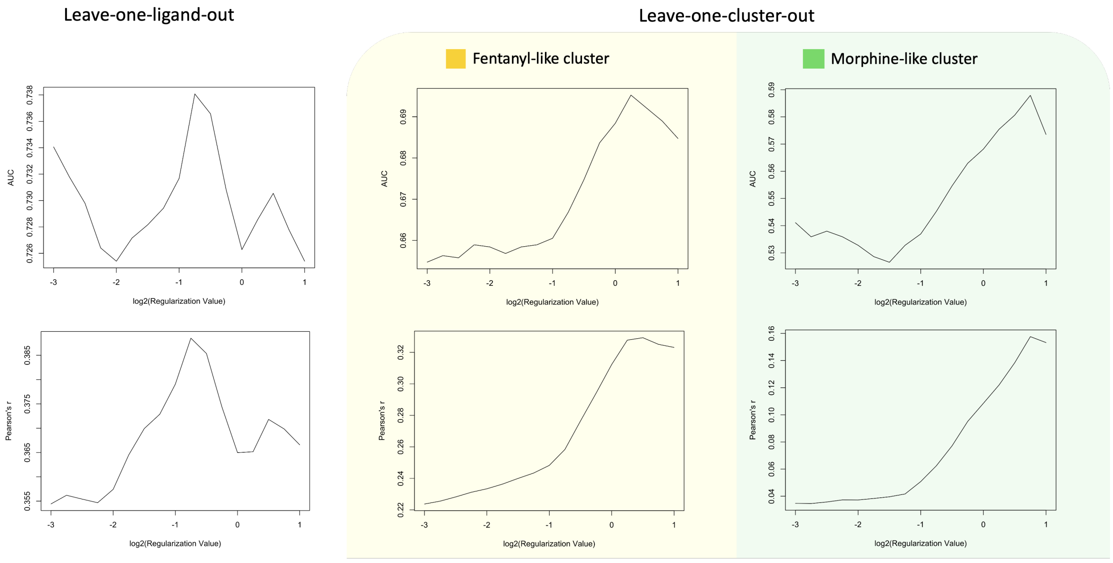


**Figure 3: Leave-one-ligand-out and leave-one-cluster-out performance (AUC and Pearson's R) over all LASSO Regularization Values ($\alpha$).**

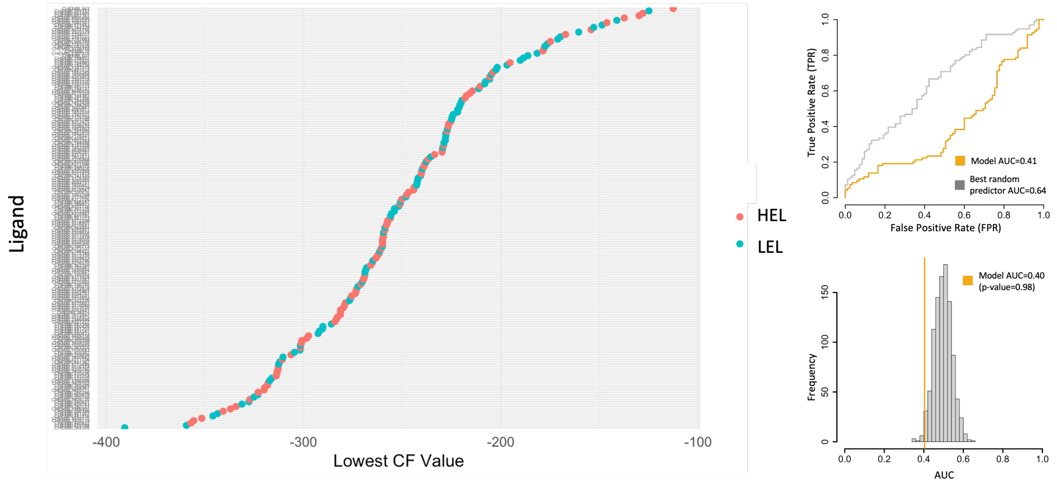


**Figure 4: (A)**Lowest CF pose per ligand in crescent order. (**B**) ROC curve against best random predictor over 1000 simulations. (**C**) AUC distribuition

### REFERENCES

[1] Vincent Frappier and Rafael J. Najmanovich. A Coarse-Grained Elastic Network Atom Contact Model and Its Use in the Simulation of Protein Dynamics and the Prediction of the Effect of Mutations. *PLoS Computational Biology*, 10(4), 2014.

[2] Francis Gaudreault and Rafael J. Najmanovich. FlexAID: Revisiting Docking on Non-Native-Complex Structures. *Journal of Chemical Information and Modeling*, 55(7):1323–1336, 7 2015.
